# Host infection selects for sRNA variants that drive bacterial social cheating

**DOI:** 10.64898/2025.12.19.695336

**Authors:** Quentin Dubois, Typhaine Brual, Marcel Sprenger, Véronique Utzinger, Romain Mercier, Jérémy Cigna, Agnès Rodrigue, Kai Papenfort, Laetitia Attaiech, Erwan Gueguen

## Abstract

Single-nucleotide mutations in regulatory small RNAs (sRNAs) represent a largely unexplored route to social cheating during bacterial infection. Here we show that *arcZ*, encoding a conserved Hfq-dependent sRNA in the plant pathogen *Dickeya solani*, undergoes recurrent single-nucleotide mutations during potato tuber infection. Five distinct *arcZ* alleles define a graded series of ArcZ functional impairment in which antifungal and protease activities are progressively reduced while virulence is compromised only in a subset of alleles. All five variants gain a fitness advantage exclusively during co-infection with wild-type *arcZ_1_* cells. This advantage is strongest when variants are rare and declines as cheater frequency increases, ultimately correlating with reduced total bacterial productivity, the canonical signature of social cheating and a tragedy of the commons. No fitness benefit is detected *in vitro*. Transcriptomic profiling of five *arcZ* variants identifies a conserved core of 186 downregulated genes enriched in secreted and diffusible functions. Systematic genetic dissection, supported by quantitative proteomics, establishes BudAB-dependent acetoin production as the dominant cooperative public good exploited during infection. Wild-type *arcZ_1_*cells divert pyruvate flux toward acetoin via the BudAB pathway, maintaining a less acidic tissue environment that sustains disease progression; *arcZ* variants, which repress this pathway, exploit the resulting pH buffering without contributing to it. These results establish pleiotropic regulatory sRNAs as a previously unrecognized class of mutational targets for the emergence of social cheaters during host infection.

**Statement:** Bacterial populations are not genetically uniform: variants that exploit the cooperative activities of their neighbors, social cheaters, can emerge rapidly within infected tissue and undermine collective bacterial productivity. We show that single-nucleotide mutations in the sRNA ArcZ of *Dickeya solani* are recurrently selected during potato infection, generating cheater variants that cease contributing to a shared metabolic benefit. Wild-type cells divert pyruvate toward acetoin, buffering tissue pH and sustaining disease; mutant variants exploit this pH maintenance without producing acetoin themselves. Because a single *arcZ* mutation simultaneously represses a conserved set of 186 genes, including genes involved in secreted and diffusible functions, regulatory sRNA genes represent a previously unrecognized and unusually efficient mutational target for the evolution of social cheating during infection.

## Introduction

In microbial communities, cooperative traits that benefit the group impose metabolic costs on individual producers, creating selective pressure for variants that reduce their own contribution while continuing to exploit shared benefits. These social cheaters gain a frequency-dependent fitness advantage. Indeed, they are favored when rare, as cooperators provide abundant public goods, but their advantage erodes as cheater frequency increases and the public good pool diminishes, ultimately generating a tragedy of the commons in which collective productivity declines (1, 2). In bacteria, cheating has been documented for a wide range of costly extracellular functions including siderophores, extracellular enzymes, quorum-sensing signals, and virulence-associated secreted factors (3–6). In all characterized cases, the emergence of cheater variants has been linked to mutations in protein-coding genes, including transcriptional regulators (7). Whether mutations in non-coding regulatory RNAs, which can control multiple cooperative outputs simultaneously through a single molecular locus, could provide a more direct route to social cheating during infection has not been explored.

*Dickeya solani* is a pectinolytic member of the *Pectobacteriaceae* and a major causal agent of soft rot and blackleg disease in potato (*Solanum tuberosum*) (8, 9). It is one of the most economically damaging bacterial phytopathogens in Europe since its emergence in the early 2000s (10, 11). Its pathogenicity relies on a suite of extracellular and diffusible molecules that act beyond the producing cell and represent candidate public goods. During infection, plant cell wall-degrading enzymes (PCWDEs), including pectinases, cellulases, and proteases, are secreted into host tissue, macerating cell walls and releasing nutrients accessible to the entire local bacterial population (11). *D. solani* also produces secondary metabolites with antimicrobial activity, including the antiascomycetal solanimycin, the phytotoxic zeamine, and the antioomycetal oocydin (12, 13). Pectinolytic bacteria including *Dickeya* and *Pectobacterium* species produce acetoin and 2,3-butanediol via the BudAB pathway. This metabolic activity has been associated with limitation of tissue acidification in a manner that sustains maceration and disease progression (14). Because acetoin-mediated pH buffering modifies the shared infection environment rather than being directly consumed by neighboring cells, it constitutes an environmentally diffuse cooperative output whose susceptibility to exploitation by non-producing variants has not been investigated.

ArcZ is a conserved Hfq-dependent sRNA across *Enterobacterales* (15, 16). It is transcribed as a ∼130-nucleotide (nt) precursor processed by RNase E to release a ∼60-nt 3’ isoform, the regulatory-active species (17). This processed form acts as a global pleiotropic post-transcriptional regulator, controlling more than 10% of the bacterial genome in model *Enterobacterales*, including master regulators of the general stress response (*rpoS*) and motility (*flhDC*) (18, 19). ArcZ also controls secondary metabolite production in the insect pathogen, *Photorhabdus laumondii* and *Xenorhabdus szentirmaii* (20). In *Dickeya dadantii*, ArcZ activates PCWDE production through direct Hfq-dependent base-pairing with the 5’ UTR of the *pecT* mRNA (21). In *D. solani*, ArcZ (59 nt) activates the production of solanimycin and zeamine (12). Because ArcZ coordinately activates multiple costly extracellular outputs, progressive reduction of its intracellular abundance or structural integrity could differentially impair downstream cooperative outputs according to their regulatory thresholds. Thus, mutational inactivation of *arcZ* could simultaneously uncouple multiple cooperative contributions.

Conflicting phenotypic descriptions of the *D. solani* reference strain IPO 2222 regarding antifungal activity across European laboratories (12, 13) raised the possibility that distinct *arcZ* alleles coexist within the founding stock, reflecting host-driven selection at this locus. Here we show that single-nucleotide mutations in *arcZ* recurrently emerge during potato tuber infection, generating variants that satisfy the canonical criteria for social cheating by exploiting BudAB-dependent acetoin production and the pH buffering it provides as a key public good. These findings identify mutations in pleiotropic regulatory sRNAs as a previously unrecognized and unusually direct route to the emergence of social cheaters during host infection.

## Results

### Convergent selection of *arcZ* variants during *D. solani* potato tuber infection

Sub-isolates recovered from the reference stock of IPO 2222 held in European collections showed markedly divergent antifungal phenotypes. Stocks from France, Belgium, Poland, and the United Kingdom failed to inhibit *Kluyveromyces lactis* growth (Sol^-^ phenotype), whereas the Scottish stock retained antifungal activity (Sol^+^ phenotype) (Fig. S1A). Sequence analysis revealed that the former stocks carry the *arcZ_2_* allele, while the Scottish stock carries the distinct *arcZ_3_* allele. (Table S1). These allelic differences were associated with differences in potato tuber maceration and altered ArcZ processing (Fig. S1B-C).

To determine whether this allelic heterogeneity originated from the founding stock, we analyzed sub-isolates recovered from the IPO 2222 reference cryotube from the Netherlands collection. Whole-genome sequencing confirmed that DS624 (*arcZ_1_*), DS623 (*arcZ_2_*), and DS625 (*arcZ_3_*) are isogenic, differing exclusively at the *arcZ* locus, with no additional SNPs detected elsewhere in their genomes. DS624 was therefore used as the *arcZ_1_* reference strain in all subsequent phenotypic and molecular analyses (Table S1). These sub-isolates, recovered as individual colonies and confirmed by whole-genome sequencing as isogenic single-allele clones, carried distinct *arcZ* alleles and displayed corresponding differences in antifungal activity (Fig. 1A). To assess the composition of the founding stock itself, amplicon sequencing was performed directly on DNA extracted from the bulk cryotube population. This revealed the coexistence of three arcZ alleles, the wild-type *arcZ_1_*, *arcZ_2_* (G90A), and *arcZ_3_* (G101T), at frequencies of approximately 2%, 91%, and 4%, respectively (Table S2). Thus, the reference cryotube is a genetically heterogeneous population from which isogenic single-allele sub-isolates can be recovered by restreaking.

**Figure 1.**
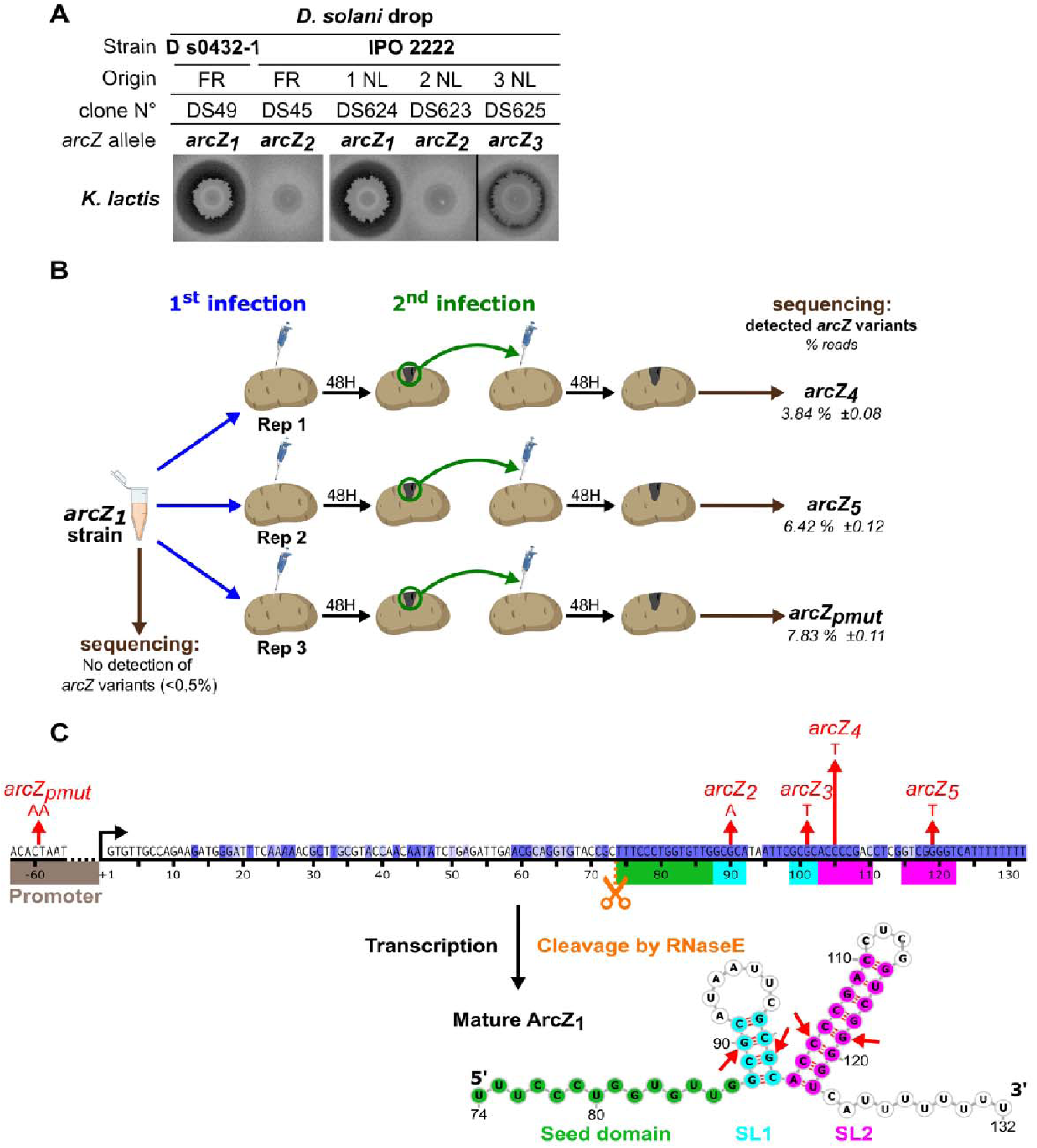
Natural coexistence and experimental emergence of *arcZ* variants in *D. solani*. **(A)** Antifungal activity against *K. lactis*. The Sol⁺ strain D s0432-1 (DS49, *arcZ_1_*) and the Sol⁻ IPO 2222 strain (DS45, *arcZ_2_*) served as positive and negative controls, respectively (12). Sub-isolates recovered from the original IPO 2222 reference cryotube from the Netherlands collection (DS624, *arcZ_1_*; DS623, *arcZ_2_*; DS625, *arcZ_3_*) are shown alongside the controls. Representative images from four biological replicates are shown. See Materials and Methods for assay conditions. **(B)** Experimental emergence of *arcZ* variants during potato tuber infection. The *arcZ_1_* strain DS624 was subjected to two successive 48 h infection cycles across three independent biological replicates. No other *arcZ* variant was detected above the 0.5% threshold in the initial inoculum. After two infection cycles, one distinct variant was recovered per replicate: *arcZ_4_* (C105T) from replicate 1, *arcZ_5_* (G119T) from replicate 2, and *arcZ_pmut_*(double promoter substitution CT➔AA at positions −60/−59, leaving the sRNA sequence unchanged) from replicate 3. Variant frequencies were quantified by Illumina amplicon sequencing of the *arcZ* locus (see Materials and Methods). Values are mean ± SD from three independent biological replicates. See Table S2 for full variant frequencies and Table S5 for strain designations. **(C)** Schematic representation of the *arcZ* locus and predicted secondary structure of ArcZ_1_. The *arcZ* locus shows the transcription start site (+1) and multiple features of the ArcZ sRNA. Sequence conservation across *arcZ* is indicated by a blue gradient at each nucleotide position (from white to dark blue, as in Fig. S7). The RNase E cleavage site is marked in orange. Sequences forming the predicted stem-loops (SL) according to RNAfold v2.0 (ViennaRNA Package 2.0) are in turquoise (SL1) and pink (SL2). The promoting region of *arcZ* is in brown. The seed sequence involved in the recognition of target mRNAs is indicated in green. The red arrows indicate the positions of mutations present in the different *arcZ* variants.

**Figure S1.**
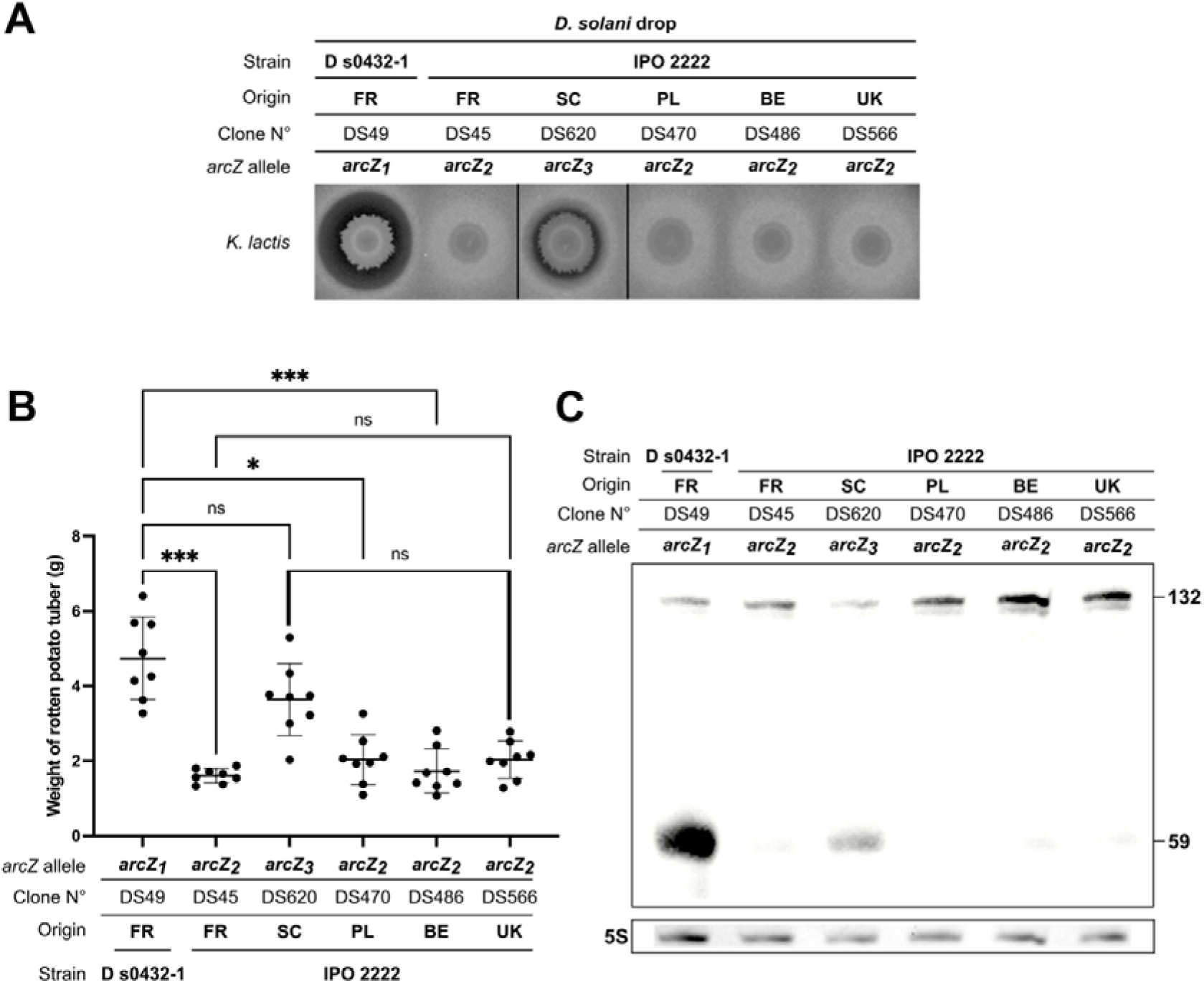
Phenotypic and molecular heterogeneity among *D. solani* stocks from European collections carrying distinct *arcZ* alleles. See Table S5 for full strain details. **(A)** Antifungal activity against *K. lactis*. The Sol⁺ strain D s0432-1 (clone DS49, *arcZ_1_*, France) and five IPO 2222-derived stocks from European collections are shown: DS45 (*arcZ_2_*, France), DS620 (*arcZ_3_*, Scotland), DS470 (*arcZ_2_*, Poland), DS486 (*arcZ_2_*, Belgium), and DS566 (*arcZ_2_*, United Kingdom). Representative images from four biological replicates are shown. **(B)** Potato tuber maceration caused by the indicated strains. Bars represent mean ± SD from eight biological replicates. Statistical comparisons were performed using pairwise Mann-Whitney tests. ***P < 0.001; *P < 0.05; ns, not significant. **(C)** Northern blot analysis of ArcZ sRNA. Strains were grown in M63 minimal medium supplemented with 1% sucrose at 30°C to OD_600_ = 0.8. The full-length precursor (132 nt) and the processed mature form (59 nt) were detected with an *arcZ*-specific probe; 5S rRNA served as a loading control. A representative blot from three independent experiments is shown.

Because IPO 2222 was originally isolated from a diseased potato plant, we next asked whether the potato tuber environment itself could promote the emergence or enrichment of *arcZ* variants. DS624 (*arcZ_1_*) was subjected to two successive 48 h infection cycles in potato tubers across three independent biological replicates (Fig. 1B). No *arcZ* variant (other than *arcZ_1_*) was detected above the 0.5% threshold in the initial inoculum. After two infection cycles, one distinct variant was recovered from each replicate: *arcZ_4_* (C105T substitution, clone DS931) from replicate 1, *arcZ_5_* (G119T substitution, clone DS944) from replicate 2, and *arcZ_pmut_* (double substitution in *arcZ* promoter-region, clone DS945) from replicate 3, at final population frequencies of 3.8%, 6.4%, and 7.8%, respectively (Fig. 1B-C, Table S2). Both *arcZ_4_* and *arcZ_5_* substitutions are in the predicted terminator stem-loop. No *arcZ* variant (other than the initial *arcZ_1_*) was detected after approximately 30 generations of *in vitro* passaging in LB medium (Table S2). Taken together, the independent recovery of distinct *arcZ* variants in each infection replicate, combined with the absence of detectable diversification *in vitro*, supports recurrent selection at the *arcZ* locus in the potato tuber environment. Whole-genome sequencing confirmed that DS931, DS944, and DS945 differ from DS624 only at the *arcZ* locus, with the single exception of a synonymous substitution in DS945 (a C to T change in A4U42_14900, a putative sugar ABC transporter, leaving the valine codon unchanged). Because this change is silent and confined to one variant, the three strains were used as a panel of near-isogenic variants for all subsequent analyses (Table S1).

### *arcZ* variants display graded defects in ArcZ accumulation, extracellular activities, and virulence-associated phenotypes

To characterize the molecular and phenotypic consequences of the five *arcZ* mutations, we compared all variants with the *arcZ_1_* reference strain across a series of functional assays. *arcZ_1_* produced a strong *K. lactis* inhibition halo, whereas *arcZ_2_*and *arcZ_5_* displayed a complete loss of antifungal activity. *arcZ_3_*, *arcZ_4_*, and *arcZ_pmut_* showed significantly reduced but detectable halos (Fig. 2A, Fig. S2A).

**Figure 2.**
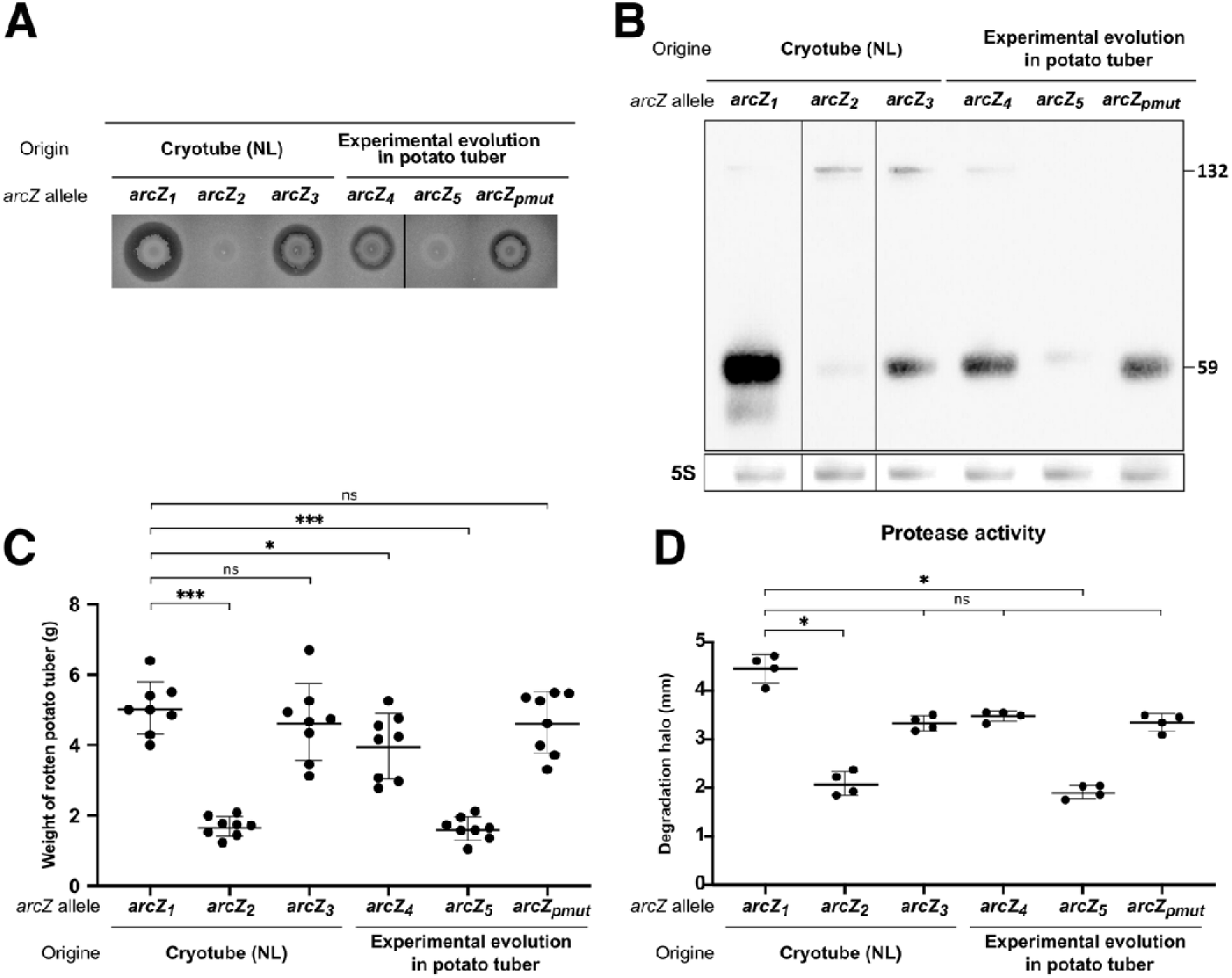
*arcZ* variants display graded defects in ArcZ accumulation, tuber maceration, and extracellular activities. All panels compare the *arcZ_1_* reference strain (DS624, Cryotube NL) with the five *arcZ* variants: DS623 (*arcZ_2_*) and DS625 (*arcZ_3_*), recovered from the same cryotube, and DS931 (*arcZ_4_*), DS944 (*arcZ_5_*), and DS945 (*arcZ_pmut_*), isolated after experimental evolution in potato tubers. See Table S5 for full strain details. **(A)** Antifungal activity against *K. lactis*. Representative images from four biological replicates are shown. Quantification is provided in Fig. S2A. **(B)** Northern blot analysis of ArcZ sRNA. The full-length precursor (132 nt) and the processed regulatory-active form (59 nt) were detected with an *arcZ*-specific probe; 5S rRNA served as a loading control. A representative blot from four independent experiments is shown; quantification is provided in Fig. S2B. **(C)** Potato tuber maceration. Bars represent mean ± SD from eight biological replicates. **(D)** Protease activity measured as degradation halo diameters on indicator plates. Bars represent mean ± SD from four biological replicates. (C and D) Statistical comparisons were performed using pairwise Mann-Whitney tests against DS624 (*arcZ_1_*). *P < 0.05; ***P < 0.001; ns, not significant.

**Figure S2.**
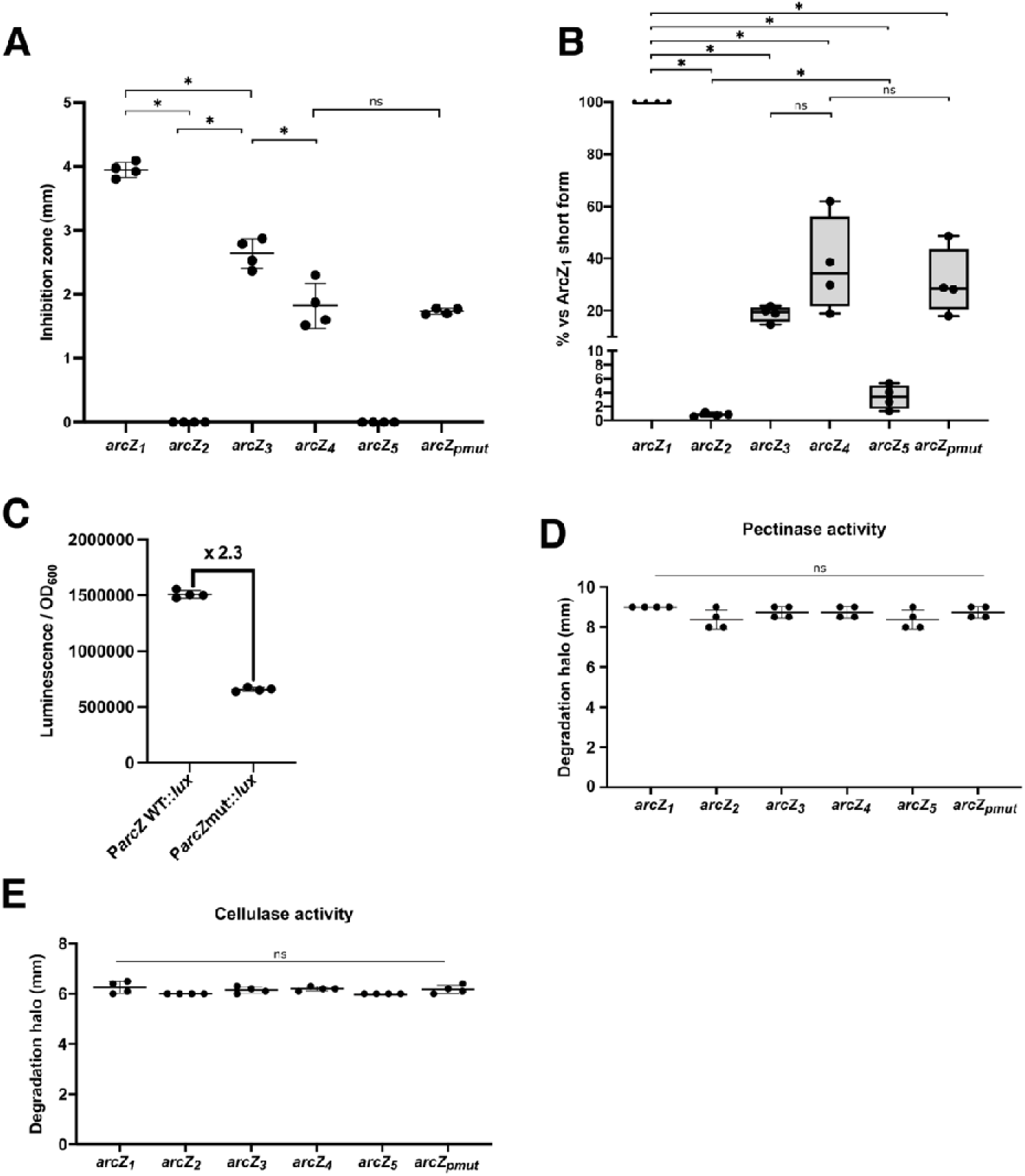
Quantification of antifungal activity, processed ArcZ accumulation, and *arcZ* promoter activity. All panels show data for the *arcZ_1_* reference strain DS624 and the five *arcZ* variants: DS623 (*arcZ_2_*) and DS625 (*arcZ_3_*), recovered from the same cryotube, and DS931 (*arcZ_4_*), DS944 (*arcZ_5_*), and DS945 (*arcZ_pmut_*), isolated after experimental evolution in potato tubers. See Table S5 for full strain details. Statistical comparisons were performed using pairwise Mann-Whitney tests. *P < 0.05; ns, not significant. **(A)** Quantification of *K. lactis* inhibition zone lengths corresponding to Fig. 2A. Inhibition zones were measured from the colony edge to the outer limit of clearing. Assay conditions as described in Fig. 2A. Bars represent mean ± SD from four biological replicates; dots represent individual replicates. **(B)** Quantification of processed ArcZ (59 nt) abundance from Northern blots corresponding to Fig. 2B. Signal intensities were normalized to 5S rRNA and expressed as a percentage of the DS624 (*arcZ*₁) mean. Note the broken y-axis between 8% and 10%. Box plots show four independent biological replicates; triangles indicate individual values. **(C)** Promoter activity of the wild-type (P*_arcZ_*_WT_) and mutant (P*_arcZ_*_mut_) *arcZ* promoter alleles measured by *lux* transcriptional fusions. The P*_arcZ_*_mut_ allele carries a dinucleotide substitution of the bases CT to AA at positions -60 and -59 relative to the transcription start site, which reduces promoter activity 2.3-fold without altering the sRNA sequence. Luminescence was measured during exponential growth (OD_600_ = 0.3) and normalized to culture density (Luminescence units/OD_600_). Bars represent mean ± SD from three biological replicates; dots represent individual replicates. **(D and E)** Extracellular enzyme activity measured as degradation halo diameters on indicator plates: pectinase (D) and cellulase (E). Bars represent mean ± SD from four biological replicates.

Northern blot analysis showed that the processed ArcZ form (59 nt) accumulated strongly in *arcZ_1_* but was absent or strongly reduced in *arcZ_2_* and *arcZ_5_* (Fig. 2B, Fig. S2B). The full-length ArcZ species (132 nt) accumulated predominantly in *arcZ_2_*, consistent with impaired RNase E processing. In contrast, *arcZ_3_*, *arcZ_4_*, and *arcZ_pmut_*retained partial accumulation of the processed form. For *arcZ_pmut_*, the promoter mutation reduced *arcZ* transcription 2.3-fold relative to wild type, as measured using Lux reporter fusions, without altering the ArcZ RNA sequence (Fig. S2C).

We next compared the pathogenicity of the *arcZ* variants using a potato tuber maceration assay. *arcZ_2_*and *arcZ_5_* caused significantly less maceration than *arcZ_1_*, whereas *arcZ_3_*and *arcZ_pmut_* retained near-wild-type *arcZ_1_* maceration and *arcZ_4_* showed a modest but significant reduction (Fig. 2C). Pectinase and cellulase activities were not significantly different across strains (Fig. S2D, E), whereas protease activity was significantly reduced in *arcZ_2_* and *arcZ_5_*and intermediate in *arcZ_3_*, *arcZ_4_*, and *arcZ_pmut_*(Fig. 2D).

Together, these data define a graded allelic series of ArcZ impairment. *arcZ_2_* and *arcZ_5_* showed the most severe defects, whereas *arcZ_3_*, *arcZ_4_*, and *arcZ_pmut_*retained partial mature ArcZ accumulation and displayed intermediate phenotypes. The *arcZ_pmut_* variant is particularly informative because it preserves the ArcZ RNA sequence, and thus its structure and processing, but reduces *arcZ* promoter activity. This is sufficient to impair antifungal and protease outputs while preserving virulence, consistent with the threshold-linear model of sRNA-mediated regulation in which targets with lower activation thresholds are preferentially affected by reduced sRNA abundance (22, 23).

Although *arcZ* variants repeatedly emerged during potato tuber infection, several displayed reduced extracellular activities and the most severe alleles caused reduced maceration in monoculture infection. These observations suggested that their selective advantage might depend on mixed infection with ArcZ-proficient *arcZ_1_* cells.

### *arcZ* variants gain a frequency-dependent fitness advantage during infection consistent with social cheating

To determine whether *arcZ* variants gain a selective advantage during potato tuber infection, we performed pairwise competition assays in which each gentamicin-resistant (Gm^R^) *arcZ* variants (DS899, DS901, DS947, DS948, DS949) was co-inoculated with the unmarked *arcZ_1_* strain DS624 as a rare population at an initial 1:100 Gm^R^:Gm^S^ ratio. To provide a complete loss-of-function control independent of the structural effects of individual point mutations, we also included the Δ*arcZ_1_* in-frame deletion mutant (DS937), constructed in the D s0432-1 *arcZ_1_* background (12) and competed against its isogenic parent DS49. Because DS937 and DS49 are isogenic except at the *arcZ* locus, any competitive advantage observed in this pair can be attributed solely to the absence of ArcZ function, without confounding effects from other genomic differences. The Δ*arcZ_1_* mutant also allows complementation experiments: restoration of *arcZ_1_* expression from the plasmid pEGL334 in DS943 was used to confirm that the competitive phenotype is specifically due to loss of ArcZ function. This setup was designed to mimic the emergence of rare variants within an ArcZ-proficient population. All five *arcZ* variants were strongly enriched after 48 h of co-infection, with log_10_ competitive index values close to 2, corresponding to an approximately 100-fold increase relative to their initial frequency (Fig. 3A). The Δ*arcZ_1_* deletion mutant showed a comparable advantage, whereas complementation with the wild-type *arcZ_1_* allele abolished this phenotype (Fig. 3A). Self-competition controls showed that the gentamicin-resistance marker imposed no detectable fitness bias (Fig. S3A). These results indicate that loss or reduction of ArcZ function is sufficient to confer a strong competitive advantage during potato tuber co-infection.

**Figure 3.**
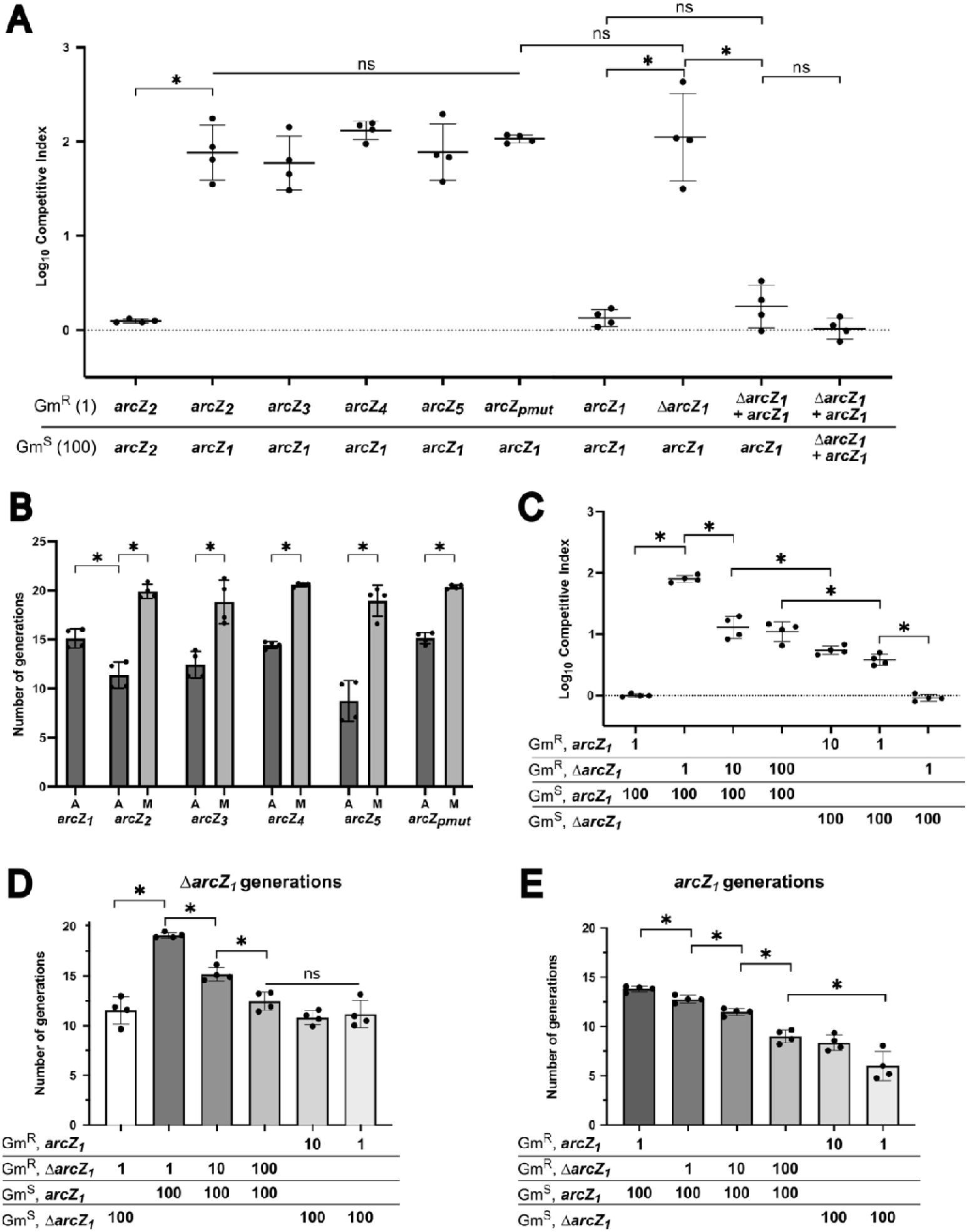
*arcZ*-deficient variants show frequency-dependent fitness advantages consistent with social cheating during potato tuber co-infection. See Table S5 for complete strain genotypes and designations. Competitive index (CI, in panels A, B and C) and number of generations (panels D and E) were calculated as described in Materials and Methods. All competition assays were performed at a total inoculum of 2 x 10^6^ bacteria per tuber; CI were calculated after 48 h. Individual results (dots) from four biological replicates with their mean ± SD are presented. Statistical comparisons were performed using pairwise Mann-Whitney tests. *P < 0.05; ns, not significant. **(A)** Competitive fitness of the *arcZ* variants and the Δ*arcZ_1_* deletion mutant during co-infection with *arcZ_1_* strain. Gentamicin-resistant (Gm^R^) strains were inoculated at a 1:100 Gm^R^:Gm^S^ ratio alongside unmarked gentamicin-sensitive (Gm^S^) competitors, as indicated below the x-axis. The Δ*arcZ_1_* deletion mutant DS937 competed against its isogenic *arcZ_1_*parent (DS49); its *arcZ_1_*-complemented derivative (DS943) served as a specificity control (12). The self-competition control (*arcZ_2_* Gm^R^ vs. *arcZ_2_* Gm^S^) confirmed the absence opf a fitness bias from the gentamicin-resistance marker. A log10 CI > 0 indicates enrichment of the Gm^R^ strain. **(B)** Number of generations completed by *arcZ* variants in single infection (A, alone, dark grey bars) and mixed infection at a 1:100 initial ratio (M, co-infection with *arcZ_1_* strain DS624, light grey bars). Generations were calculated from strain-specific CFU counts for each variant population. **(C)** Frequency-dependence of the Δ*arcZ_1_* competitive advantage. Gm^R^ and Gm^S^ derivatives of *arcZ_1_* (DS49) and Δ*arcZ_1_* (DS937) strains were competed at the initial ratios indicated below the x-axis. **(D-E)** Number of generations completed by the Δ*arcZ_1_*(D) and *arcZ_1_* (E) strains at the initial inoculum ratios indicated below the x-axis. Panels D and E are derived from the same experiments as panel C. Bar shading represents the initial *arcZ_1_*:Δ*arcZ_1_* ratio, with darker bars indicating a higher proportion of *arcZ_1_* and lighter bars indicating a higher proportion of Δ*arcZ_1_*.

**Figure S3.**
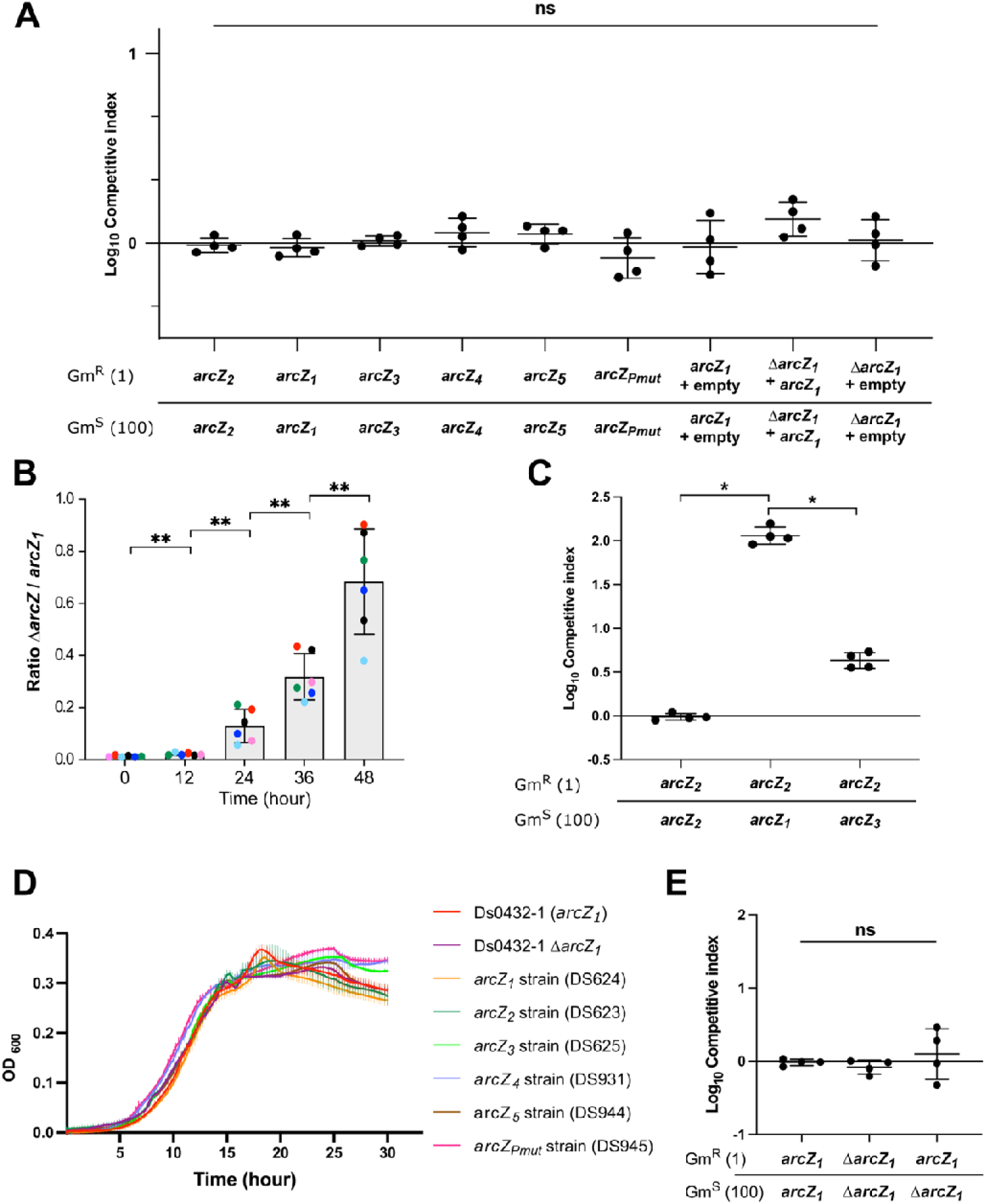
Marker controls, infection dynamics, and *in vitro* growth assays supporting the competitive advantage of *arcZ* variants. See Table S5 for full strain details. Competitive Index (CI; A, C, E) and number of generations (B) were calculated as described in Materials and Methods; data represent mean ± SD of four biological replicates (individual data points shown). Statistical comparisons were performed using pairwise Mann-Whitney tests for panels A, B, C and E as indicated. *P < 0.05; **P < 0.01; ns, not significant. **(A)** Self-competition controls confirming the absence of a fitness bias introduced by the gentamicin-resistance marker. Each Gm^R^ strain was mixed with its isogenic unmarked Gm^S^ counterpart at a 1:100 ratio (2 × 10J bacteria per tuber) and CI were calculated after 48 h. All *arcZ* variants, the Δ*arcZ_1_* deletion mutant, and the complemented derivatives were tested. **(B)** Time-course of Δ*arcZ_1_* strain (DS937) enrichment during co-infection with *arcZ_1_* strain (DS49) at an initial 1:100 ratio. CFUs of Δ*arcZ_1_*and *arcZ_1_* strains were determined at 0, 12, 24, 36, and 48 h post-inoculation in six independent biological replicates. The Δ*arcZ_1_*:*arcZ_1_*ratio of each replicate is show in a different color. The mean ± SD at each time point is presented. **(C)** Competitive fitness of *arcZ_2_* strain (DS899, Gm^R^) against *arcZ_1_*strain (DS624) or the partially impaired *arcZ_3_* strain (DS625) as Gm^S^ competitors, at a 1:100 initial ratio. This experiment tests whether the cheating advantage of a severely impaired variant depends on the level of ArcZ function in the co-infecting population. **(D)** *In vitro* growth curves of *D. solani* strains in M63 minimal medium supplemented with 0.2% glycerol at 30°C. OD_600_ was monitored every 15 min for 30 h. Curves show mean ± SD from four biological replicates. Doubling times were calculated during the exponential growth phase and compared between strains using pairwise Mann-Whitney tests. **(E)** *In vitro* competitive fitness. Gm^R^ and Gm^S^ strains were mixed at a 1:100 ratio and CI were calculated after 24 h in M63 minimal medium supplemented with 0.2% glycerol at 30°C. The 24 h duration was chosen to match the approximate number of generations completed during the 48 h *in vivo* assays.

**Figures S4:**
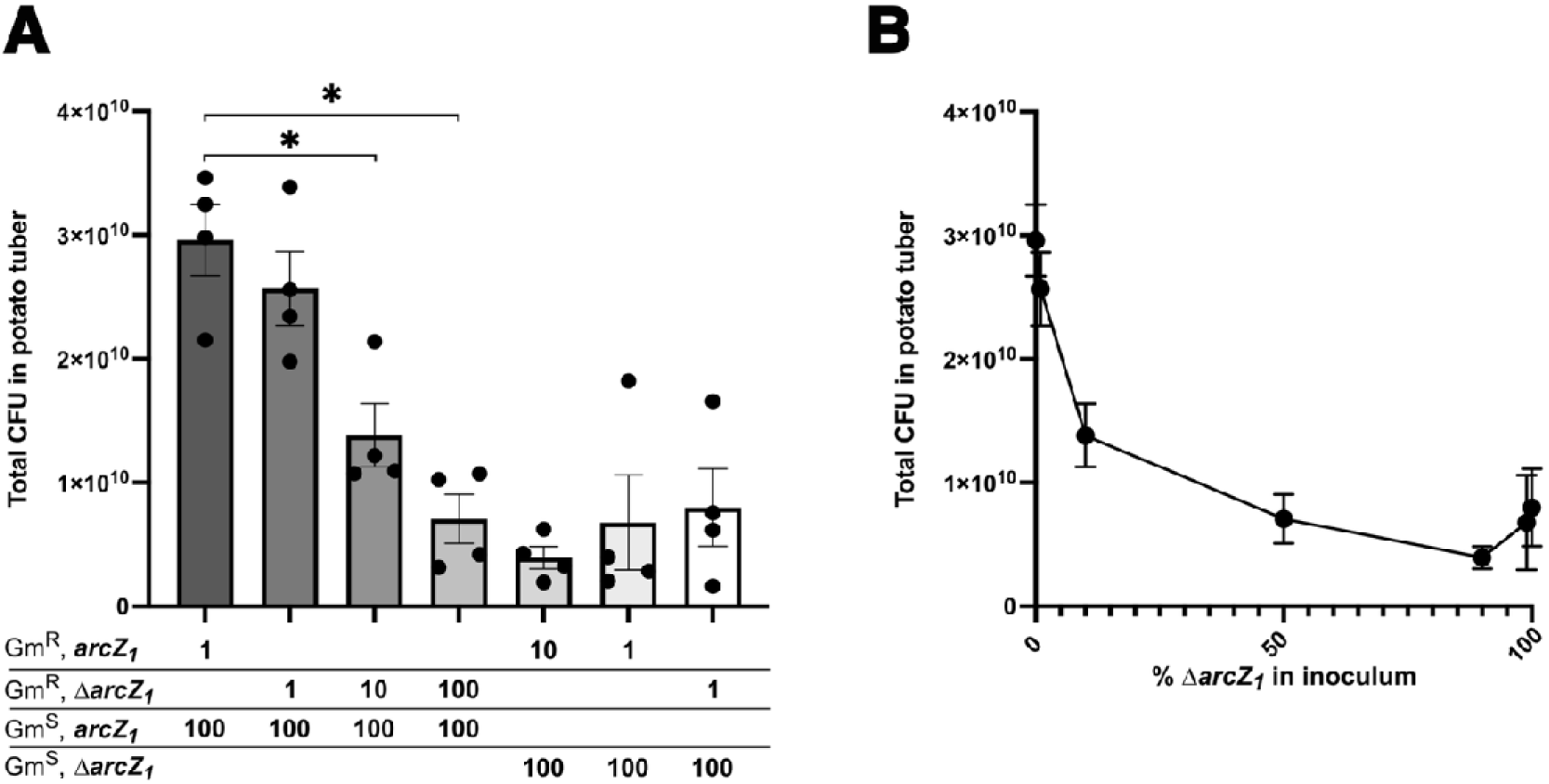
Total bacterial population size during co-infection decreases with increasing proportion of Δ*arcZ_1_* in the inoculum. All assays were performed at a total inoculum of 2 x 10^6^ bacteria per tuber; total CFUs were calculated as the sum of Gm^R^ and Gm^S^ populations recovered after 48 h. See Materials and Methods for strain details and CFU quantification. **(A)** Total CFUs recovered from potato tubers after 48 h of co-infection at the initial inoculation ratios of Gm^R^ and Gm^S^ derivatives of *arcZ_1_* (DS936) and Δ*arcZ_1_* (DS937, DS354) strains indicated below the x-axis. Bar shading represents the initial *arcZ_1_*:Δ*arcZ_1_* ratio, with darker bars indicating a higher proportion of *arcZ_1_*and lighter bars indicating a higher proportion of Δ*arcZ_1_*. Dots represent four biological replicates; bars represent mean ± SD. Statistical comparisons were performed using pairwise Mann-Whitney tests against the *arcZ_1_* monoculture condition (first bar on the left). *P < 0.05. **(B)** Total CFU recovered as a function of the initial percentage of Δ*arcZ_1_* strain in the inoculum. Data points represent mean ± SD from four biological replicates (same experiments as in panel A).

We next asked whether this enrichment reflected increased proliferation of *arcZ* variants in the presence of the wild-type *arcZ_1_* population. In single-strain infections, *arcZ_2_* and *arcZ_5_* strains, which showed the strongest defects in tuber maceration, completed fewer generations than the *arcZ_1_* strain, whereas *arcZ_3_*, *arcZ_4_*, and *arcZ_pmut_* strains proliferated at levels closer to the reference strain. In contrast, all five *arcZ* variants completed significantly more generations during mixed infection with the *arcZ_1_* strain than during single-strain infection (Fig. 3B). Growth analysis in M63 minimal medium supplemented with 0.2% glycerol revealed modest differences among strains: the *arcZ* variants and the Δ*arcZ_1_* displayed slightly shorter doubling times in exponential phase than *arcZ_1_*, although their growth curves remained largely overlapping (Fig. S3D). By contrast, the Δ*arcZ_1_* strain displayed no advantage over the *arcZ_1_* strain in pairwise competition (Fig. S3E). Thus, the competitive success of *arcZ* variants is not explained by an intrinsic growth advantage, but by a proliferation benefit specific to the potato tuber co-infection context.

To determine whether enrichment occurred progressively during infection, we monitored the Δ*arcZ_1_*strain dynamics over time during co-infection with the *arcZ_1_* strain. Starting from a 1:100 Δ*arcZ_1_*:*arcZ_1_*ratio, the relative abundance of Δ*arcZ_1_* increased from 12 h onward and continued to rise up to 48 h post-inoculation (Fig. S3B). This temporal pattern indicates that ArcZ-deficient cells are progressively enriched during infection rather than being detected only as an endpoint outcome.

We then tested whether the benefit gained by ArcZ-deficient cells depended on the functional capacity of the surrounding population. When the *arcZ_2_* strain was competed against the *arcZ_1_*strain, it displayed a strong competitive advantage, with a log_10_ CI close to 2. Interestingly, when competed against the partially impaired *arcZ_3_* variant, this advantage was significantly reduced, although it remained above zero (Fig. S3C). This reduction suggests that the benefit gained by severely impaired *arcZ* variants depends, at least in part, on the cooperative output provided by the competing population.

To test whether this advantage was frequency-dependent, we varied the initial proportion of the Δ*arcZ_1_*strain relative to the *arcZ_1_* strain. The competitive advantage of the Δ*arcZ_1_*strain was highest when the mutant was initially rare and progressively decreased as its initial frequency increased (Fig. 3C). Consistently, the Δ*arcZ_1_*strain completed the highest number of generations when rare in the presence of the *arcZ_1_* strain, and this proliferation benefit declined progressively as its proportion increased, becoming undetectable at equal inoculum ratios (Fig. 3D). Conversely, the number of generations completed by the *arcZ_1_*strain decreased as the initial proportion of the Δ*arcZ_1_* strain increased (Fig. 3E). At the population level, total CFUs recovered from tubers after 48 h infection also declined as the initial proportion of Δ*arcZ_1_* strain increased, dropping from approximately 3 x 10^10^ CFUs in *arcZ_1_* monoculture (i.e. with 0% of Δ*arcZ_1_*strain) to approximately 4 x 10^9^ CFUs when the Δ*arcZ_1_* strain was predominant (Fig. S4A-B).

Together, these results show that ArcZ-deficient variants gain a frequency-dependent advantage during potato tuber infection while reducing collective population productivity when they become abundant, a pattern consistent with social cheating.

### *arcZ* variants converge on downregulation of candidate public good associated functions

To identify molecular changes that could underlie the cheating phenotype, we performed transcriptomic profilings of the five *arcZ* variants DS623 (*arcZ_2_*), DS625 (*arcZ_3_*), DS931 (*arcZ_4_*), DS944 (*arcZ_5_*), and DS945 (*arcZ_pmut_*) relative to the *arcZ_1_* strain DS624. Using a threshold of log_2_ FC ≤ -1 and FDR-adjusted P < 0.05, we identified a core set of 186 genes consistently downregulated across all five variants (Fig. 4A). In contrast, upregulated genes were fewer and more divergent across variants, with only 13 genes induced in all five variants (Fig. S5A). This conserved downregulation signature suggests that the selective advantage of *arcZ* variants is more likely associated with coordinated repression of an ArcZ-dependent expression program than with acquisition of variant-specific functions.

**Figure 4.**
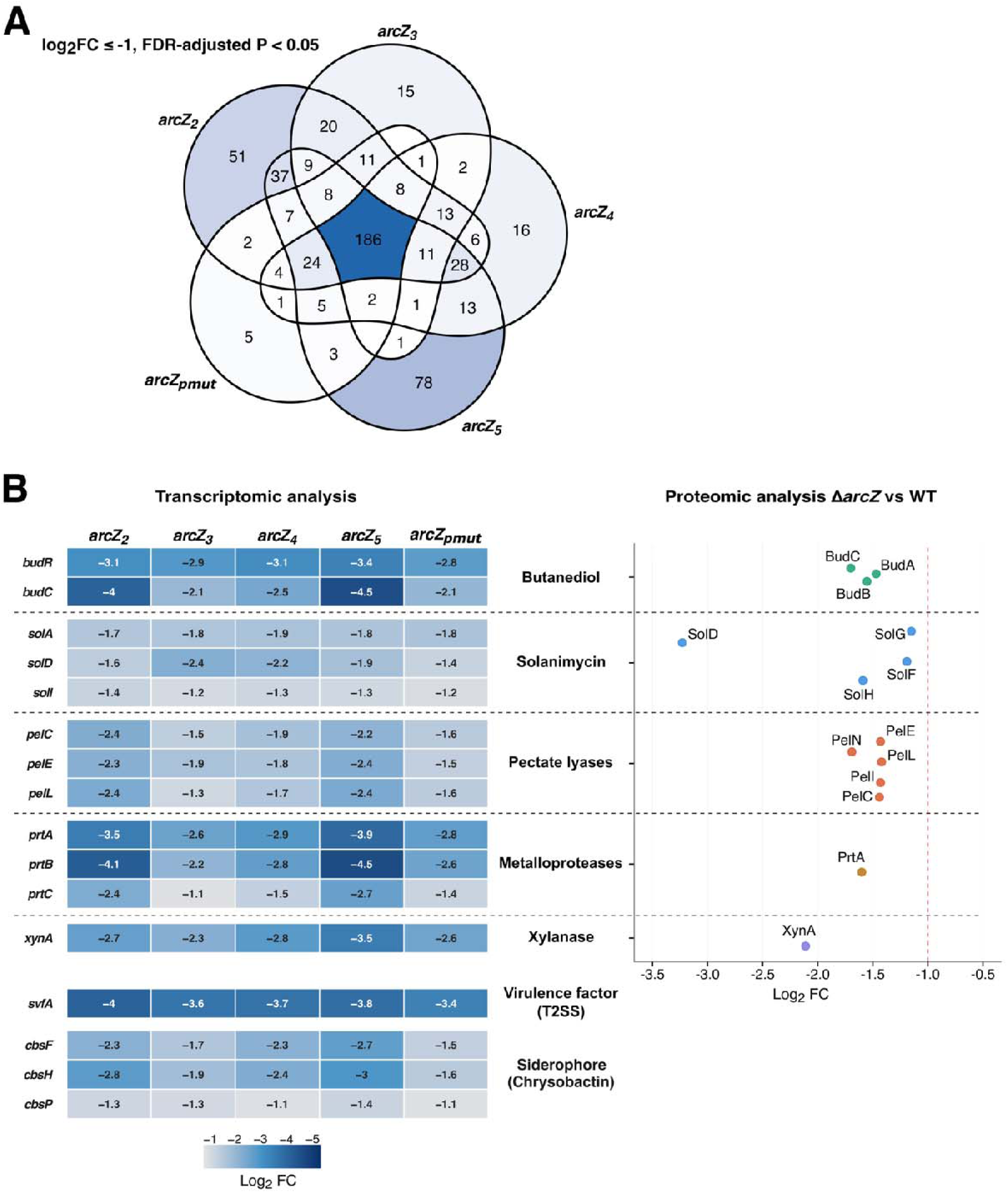
*arcZ* variants share a conserved downregulated transcriptional signature enriched in candidate public good associated functions. **>(A)** Venn diagram of genes identified by RNA-seq transcriptomic profiling as downregulated across the five *arcZ* variants relative to the *arcZ_1_* reference strain DS624 (log_2_ FC ≤ -1, FDR-adjusted P < 0.05). Strains were grown in LB medium at 30°C to OD_600_ = 0.5. The central intersection represents 186 genes downregulated in all five variants. Data are from three biological replicates per strain. **(B)** Convergent transcriptomic and proteomic evidence for downregulation of candidate public good associated functions. The two panels were generated from distinct biological samples grown under different conditions. *Left*, heatmap of log_2_ FC values for selected genes belonging to the 186-gene core downregulation set, across the five *arcZ* variants (DS623, DS625, DS931, DS944, DS945) grown in LB medium at OD_600_ = 0.5 as described in panel A. Genes are grouped by functional category; color scale indicates log_2_ FC intensity. *Right*, quantitative proteomic analysis of the Δ*arcZ_1_* deletion mutant (DS354) versus the *arcZ_1_*strain DS49, cultivated in M63 minimal medium supplemented with 1% sucrose at 30°C to OD_600_ = 0.3. The Δ*arcZ_1_*deletion mutant was used to maximize the dynamic range of ArcZ-dependent protein changes independently of the structural effects of individual point mutations present in the *arcZ* variants. Each point represents a differentially abundant protein (log_2_ FC ≤ -1, FDR-adjusted P < 0.05) assigned to the indicated functional category; the vertical dashed line indicates log_2_ FC = -1. Proteomic data are from three biological replicates. Statistical analysis for transcriptomic data: differential gene expression was assessed using the edgeR algorithm with TMM normalization and an FDR-adjusted p-value threshold of 0.05, as described above, as implemented in CLC Genomics Workbench. Statistical analysis for proteomic data: differential protein abundance was assessed by ANOVA with an FDR-adjusted p-value threshold of 0.05 using Proteome Discoverer 2.5.

**Figure S5.**
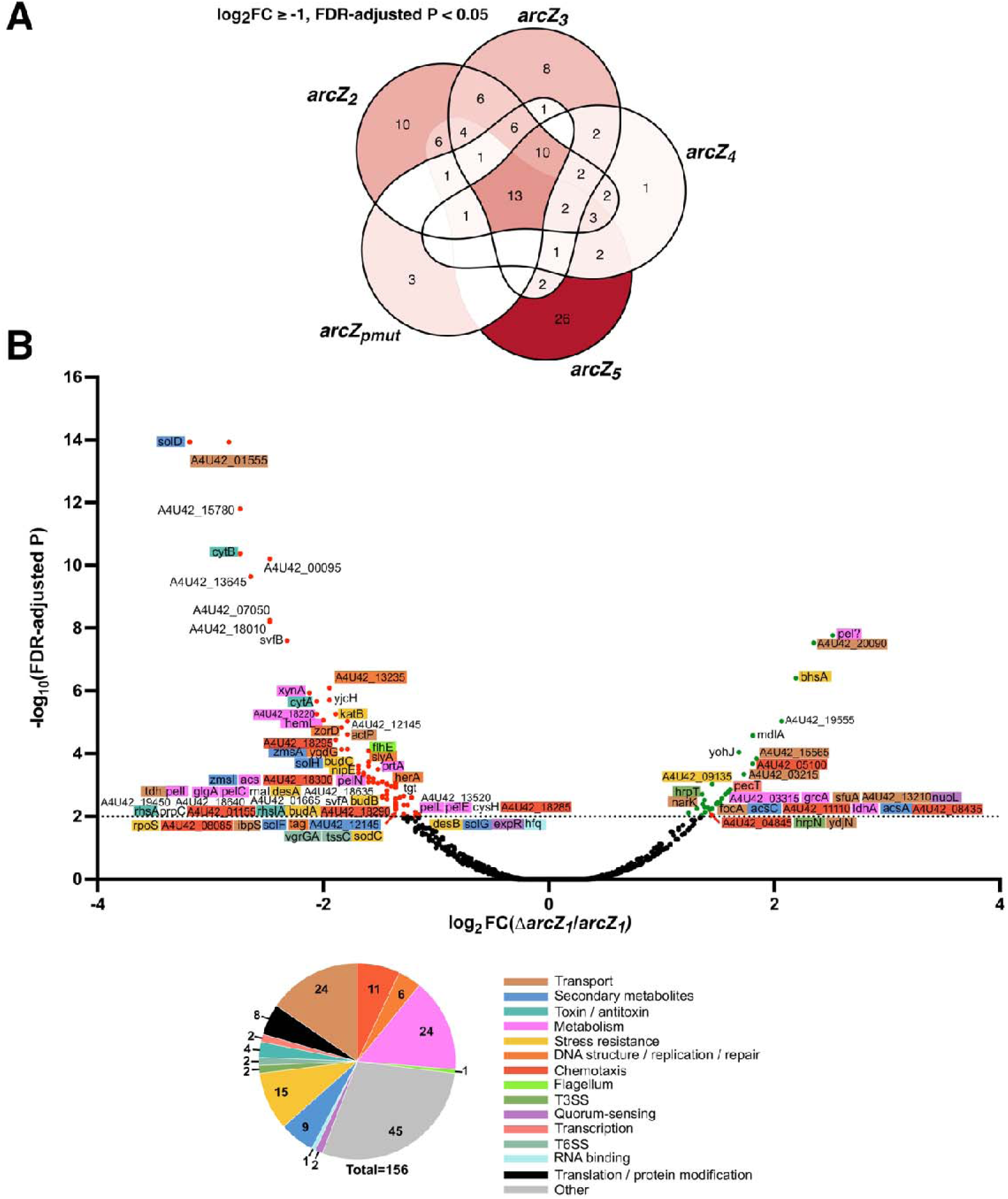
Upregulated genes in *arcZ* variants and proteomic landscape of the Δ*arcZ_1_* mutant. Note: panels A and B were generated from distinct biological samples grown under different conditions (see Materials and Methods and Fig. 4B legend). **(A)** Venn diagram of genes identified by RNA-seq transcriptomic profiling as upregulated across the five *arcZ* variants (DS623, DS625, DS931, DS944, DS945) relative to the *arcZ_1_*reference strain DS624 grown in LB medium to OD_600_ = 0.5 at 30°C (log_2_ FC ≥ 1, FDR-adjusted P < 0.05). Only 13 genes were induced in all five variants, contrasting with the 186-gene conserved downregulation core shown in Fig. 4A. Data are from three biological replicates per strain. **(B)** Volcano plot of quantitative proteomic analysis comparing the Δ*arcZ_1_* (DS354) and *arcZ_1_* strains (DS49) grown in M63 minimal medium supplemented with 1% sucrose, at 30°C, up to OD_600_ = 0.3). The x-axis shows the log_2_ protein abundance ratio (DS354/DS49); the y-axis shows -log_10_ adjusted P-value. The dotted line indicates FDR-adjusted P = 0.05. Differentially abundant proteins are labeled and color-coded by functional category (see inset). A total of 156 proteins showed significantly altered abundance (106 decreased and 50 increased in DS354 versus DS49). The pie chart summarizes the functional distribution of all 156 proteins. Data are from three biological replicates.

Functional annotation of the 186 downregulated-gene core identified several pathways encoding, regulating, or delivering extracellular and diffusible products, including the acetoin/2,3-butanediol pathway (*budC* and *budR*), solanimycin biosynthesis, pectate lyases, metalloproteases, xylanase, a type II secretion-associated virulence factor, and chrysobactin siderophore biosynthesis (Fig. 4B, left; Table S3). Many of these functions are secreted, diffusible, or expected to benefit the bacterial population during host colonization, making them candidate public good associated functions that could be exploited by non-producing variants.

To provide an independent line of evidence at the protein level, we performed quantitative proteomics on the Δ*arcZ_1_* deletion mutant DS354 relative to the *arcZ_1_* strain DS49, both cultivated in M63 minimal medium supplemented with 1% sucrose at 30°C to OD_600_ = 0.3. The Δ*arcZ_1_* deletion mutant was chosen to maximize the dynamic range of ArcZ-dependent protein changes independently of the structural effects of individual point mutations present in the *arcZ* variants. Despite the difference in biological samples and growth conditions between the two omics approaches, reduced protein abundance was observed in the same functional categories as the in the transcriptomic analysis, including acetoin/2,3-butanediol pathway enzymes, solanimycin-associated proteins, pectate lyases, metalloproteases, and xylanase (Fig. 4B, right; Fig. S5B; Table S4).

Together, these data identify a conserved ArcZ-dependent expression program enriched in candidate public good associated functions. This provided a molecular basis to test which repressed pathways contribute to the frequency-dependent advantage of *arcZ* variants during potato tuber infection.

### BudAB-dependent acetoin production is a major cooperative function exploited by *arcZ* cheaters

To identify which ArcZ-dependent functions contribute to the cheating advantage, we generated deletion mutants in the D s0432-1 background affecting major candidate public good associated pathways and measured their competitive fitness against the *arcZ_1_* strain DS49. Gm^R^-marked derivatives were used for all competition assays (Table S5). Deletion of the solanimycin cluster, chrysobactin biosynthesis genes, or the combined Δ*pelBCZ* Δ*prtBCA* mutant conferred no detectable competitive advantage against the *arcZ_1_* strain (Fig. 5A). In contrast, deletion of *budR*, the transcriptional activator of the acetoin/2,3-butanediol pathway, conferred a significant partial advantage, with a log_10_ CI of approximately 1.2 (Fig. 5A).

**Figure 5:**
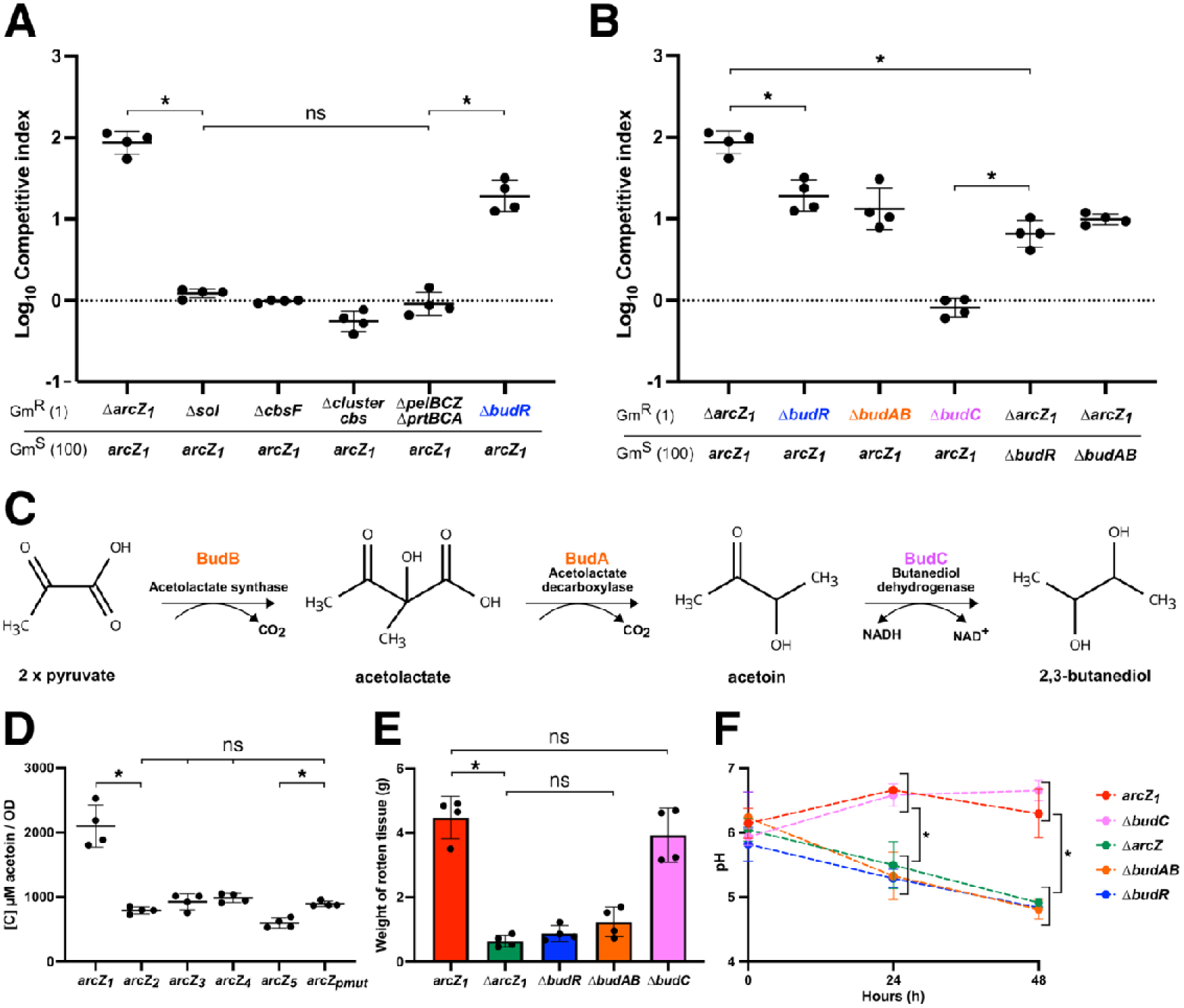
BudAB-dependent acetoin pathway activity is a major cooperative function exploited by *arcZ* cheaters. See Table S5 for full strain details. All competition assays in panels A and B were performed at an initial 1:100 Gm^R^:Gm^S^ ratio (2 x 10^6^ bacteria per tuber); Competitive Indices (CI) were calculated after 48 h as described in Materials and Methods. Color coding is used to highlight the tested components from the Bud pathway: *budR* (which esncodes the transcriptional regulator of the Bud pathway) is show in blue*, budAB*/BudA-B shown in orange and *budC*/BudC in magenta. In A, B, D and E, results of four biological replicates are presented with their mean ± SD. Statistical comparisons were performed using pairwise Mann-Whitney tests. *P < 0.05; ns, not significant. **(A)** Competitive fitness of deletion mutants targeting candidate public good associated pathways during co-infection with the *arcZ_1_* strain (DS49, Gm^S^). Gm^R^ strains tested: Δ*arcZ_1_* (DS937), Δ*sol* (DS1058), Δ*cbsF* (DS1039), Δ*cbs* cluster (DS1040), Δ*pelBCZ* Δ*prtBCA* (DS1042), and Δ*budR* (DS1003). **(B)** Competitive fitness of the Δ*arcZ_1_* and acetoin/2,3-butanediol pathway mutants against the indicated Gm^S^ competitors. The first four conditions test whether *bud* pathway deletions confer a competitive advantage against the *arcZ_1_* strain. The last two conditions test whether the competitive advantage of the Δ*arcZ_1_* strain is reduced when the cooperating population lacks BudAB-dependent activity. **(C)** Schematic of the acetoin/2,3-butanediol biosynthetic pathway. BudB (acetolactate synthase) condenses two pyruvate molecules into acetolactate; BudA (acetolactate decarboxylase) converts acetolactate into acetoin with CO_2_ release; BudC (butanediol dehydrogenase) reduces acetoin to 2,3-butanediol using NADH as cofactor. **(D)** Acetoin production by the *arcZ_1_* reference strain (DS624) and the five *arcZ* variants (DS623, DS625, DS931, DS944, DS945), measured in culture supernatants and normalized to OD_600_. **(E)** Potato tuber maceration by the *arcZ_1_* strain and the acetoin/2,3-butanediol pathway mutants in monoculture. **(F)** Tissue pH dynamics during infection at 0, 24, and 48 h post-inoculation for the indicated strains. Data represent mean ± SD from four biological replicates. Brackets indicate statistical comparisons performed using pairwise Mann-Whitney tests. *P < 0.05.

**Figure S6.**
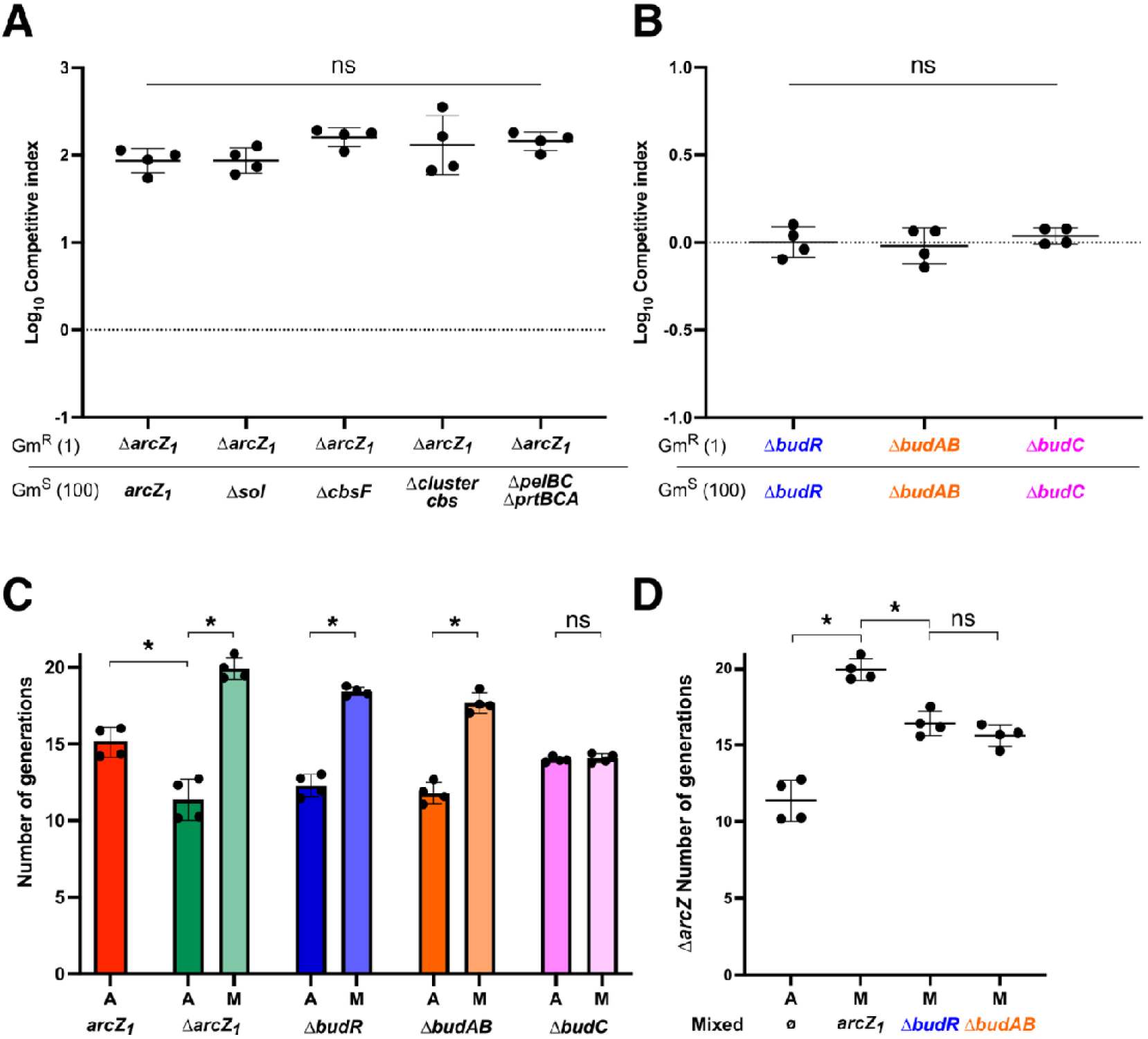
Contribution of candidate cooperative pathways to the competitive advantage and proliferation of the Δ*arcZ_1_* strain during potato tuber infection. See Table S5 for full strain details. All competition assays were performed at 1:100 Gm^R^:Gm^S^ (2 x 10^6^ bacteria per tuber) with Competitive Indices (CI) and number of generations were calculated after 48 h as described in Materials and Methods. Color coding is used to highlight the tested Bud pathway components: *budR* (which encodes the transcriptional regulator of the Bud pathway) is show in blue*, budAB*/BudA-B are shown in orange and *budC*/BudC in magenta. In A, B, C and D, results of four biological replicates are presented with their mean ± SD. Statistical comparisons were performed using pairwise Mann-Whitney tests. *P < 0.05; ns, not significant. **(A)** Self-competition controls for deletion mutants used in Fig. 5A (Δ*arcZ_1_*, Δ*sol*, Δ*cbsF*, Δ*cbs* cluster, Δ*pelBCZ* Δ*prtBCA*). **(B)** Self-competition controls for acetoin/2,3-butanediol pathway mutants used in Fig. 5B (Δ*budR*, Δ*budAB*, Δ*budC*). **(C)** Number of generations completed by the *arcZ_1_*, Δ*arcZ_1_*, Δ*budR*, Δ*budAB*, and Δ*budC* strains in single infection (A, alone, dark colored bars) and mixed infection at a 1:100 initial ratio (M, co-infection with the *arcZ_1_*strain DS49, light colored bars). Generations were calculated from strain-specific CFU counts as described in Materials and Methods. **(D)** Number of generations completed by the Δ*arcZ_1_* strain (DS937) during co-infection at a 1:100 initial ratio with *arcZ_1_* (DS49), Δ*budR* (DS883), or Δ*budAB* (DS708) strains, or in monoculture. This panel tests whether the proliferation benefit of the Δ*arcZ_1_* strain depends on BudAB-dependent activity in the surrounding population.

**Figure S7.**
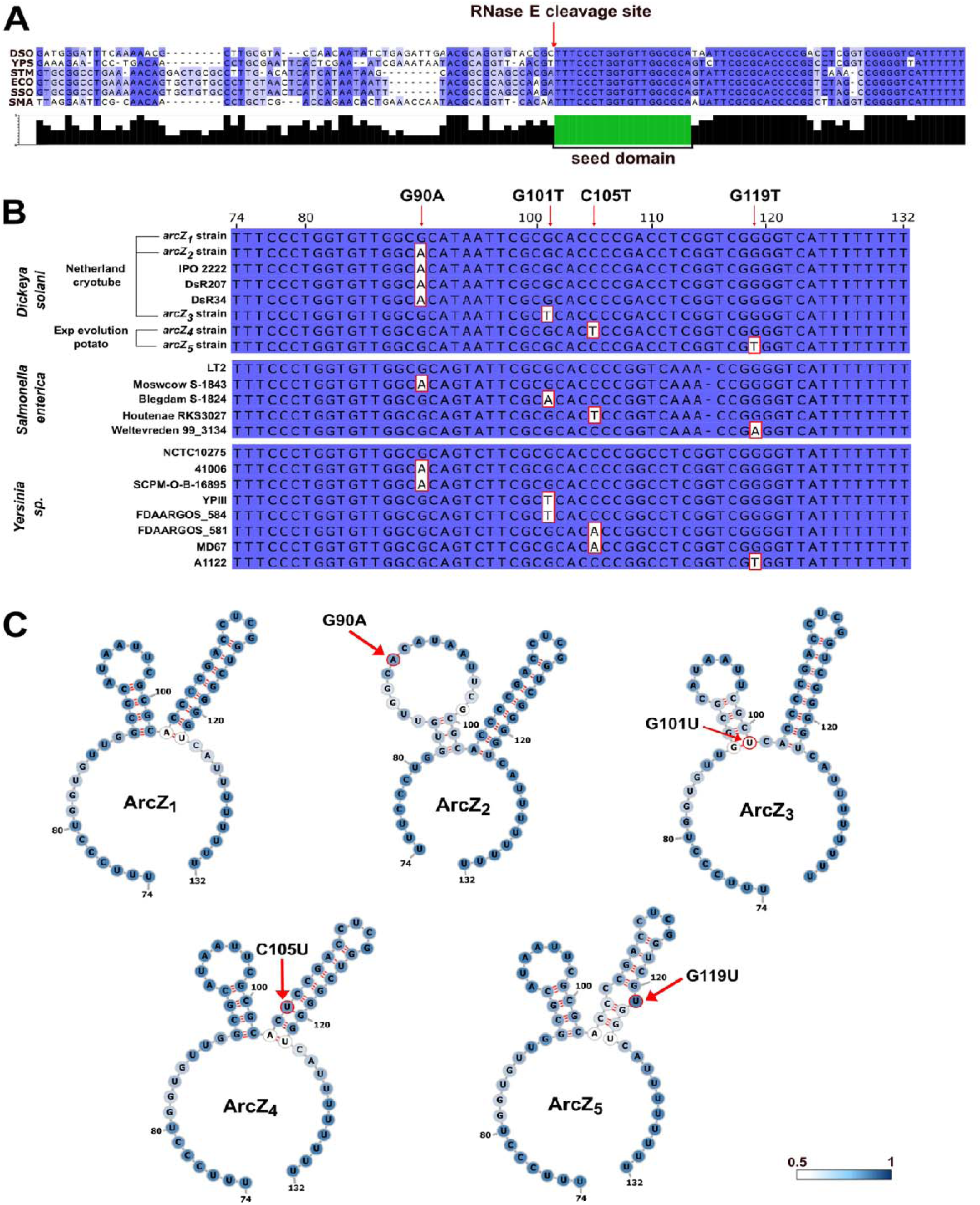
*arcZ* mutations identified in *D. solani* affect positions that are naturally variable across *Salmonella enterica* and *Yersinia sp*. Isolates. **(A)** Multiple sequence alignment of ArcZ sRNA homologs from representative *Enterobacterales*: *D. solani* (DSO), *Yersinia* sp. (YPS), *Salmonella enterica* (STM), *Escherichia coli* (ECO), *Shigella sonnei* (SSO), and *Serratia marcescens* (SMA). The predicted RNase E cleavage site is indicated above the alignment; the seed region of the processed form is highlighted in red in the conservation histogram below. Alignment performed with ClustalW and visualized with Jalview v2.11.5.1. **(B)** Nucleotide alignment of the processed ArcZ sRNA (59 nt) from *D. solani* isolates and variants, *S. enterica*, and *Yersinia* sp. strains. Red arrows above the alignment indicate the four positions mutated in *D. solani* variants (G90A, G101T, C105T, G119T); polymorphic positions are shown on a white background. The same substitutions occur at equivalent positions in naturally occurring *S. enterica* and *Yersinia* sp. isolates, indicating that these positions are structurally tolerant of variation across *Enterobacterales*. Alignment performed with ClustalW and visualized with Jalview v2.11.5.1. Sequence accession numbers are provided in Table S7. **(C)** Predicted secondary structures of the processed ArcZ sRNA (59 nt) from *arcZ_1_* through *arcZ_5_* strains. The blue gradient indicates base-pairing probability (0.5-1.0); red arrows indicate substitution positions. The *arcZ_pmut_* promoter variant leaves the sRNA sequence unchanged and was therefore not modeled separately. Structures predicted using RNAfold v2.0 (ViennaRNA Package 2.0).

**Figure S8.**
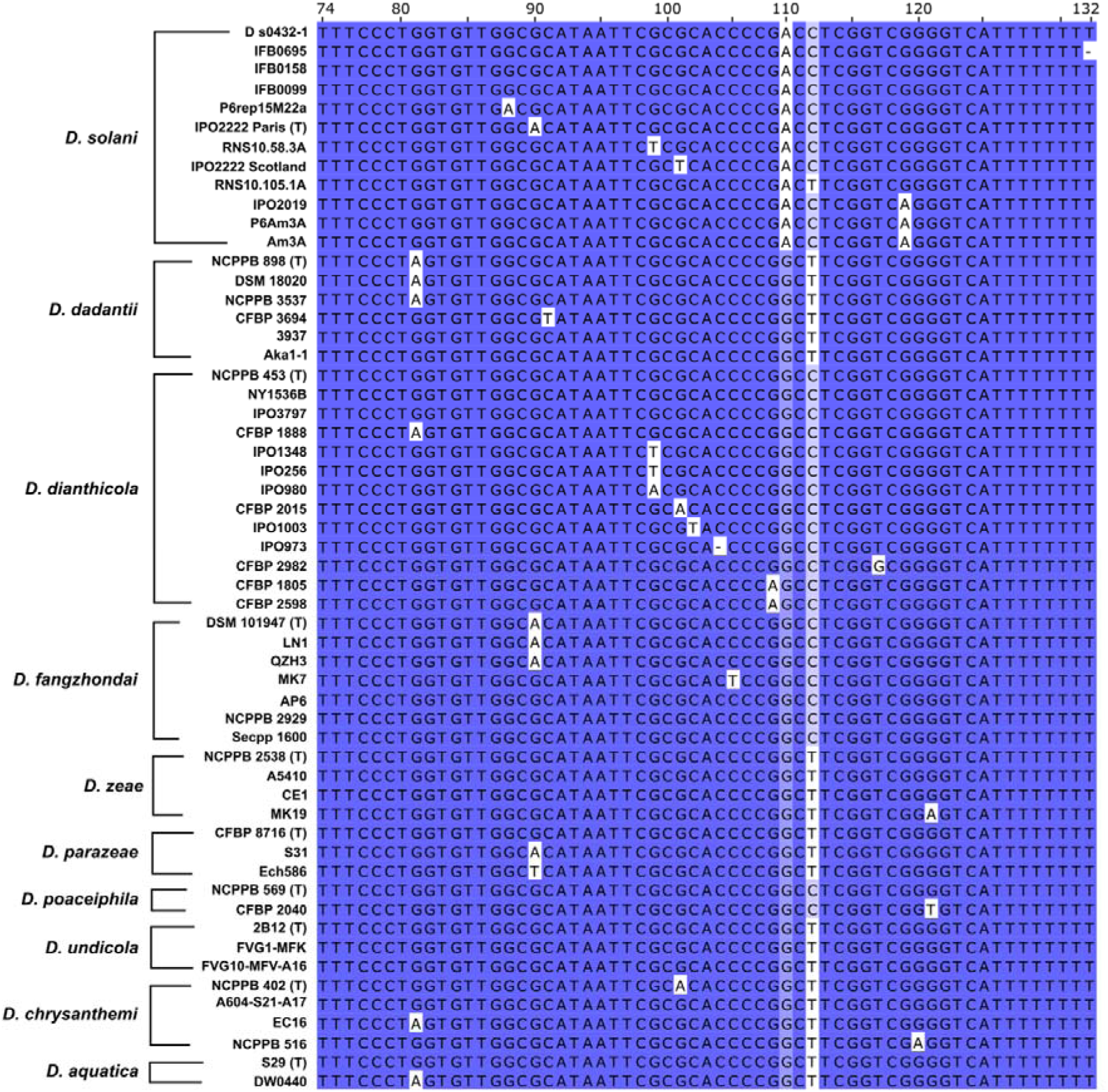
Conservation of the processed ArcZ sequence across the *Dickeya* genus. Multiple sequence alignment of the *arcZ* genomic region encoding the processed sRNA form (∼59 nt) from representative strains of ten *Dickeya* species, organized by species phylogeny. For each species, the reference sequence is listed first; type strains are indicated with (T). Dark blue shading indicates nucleotide identity; polymorphic positions are shown on a white background. Alignment performed with ClustalW and visualized with Jalview v2.11.5.1. Sequence accession numbers are provided in Table S7.

**Figure S9.**
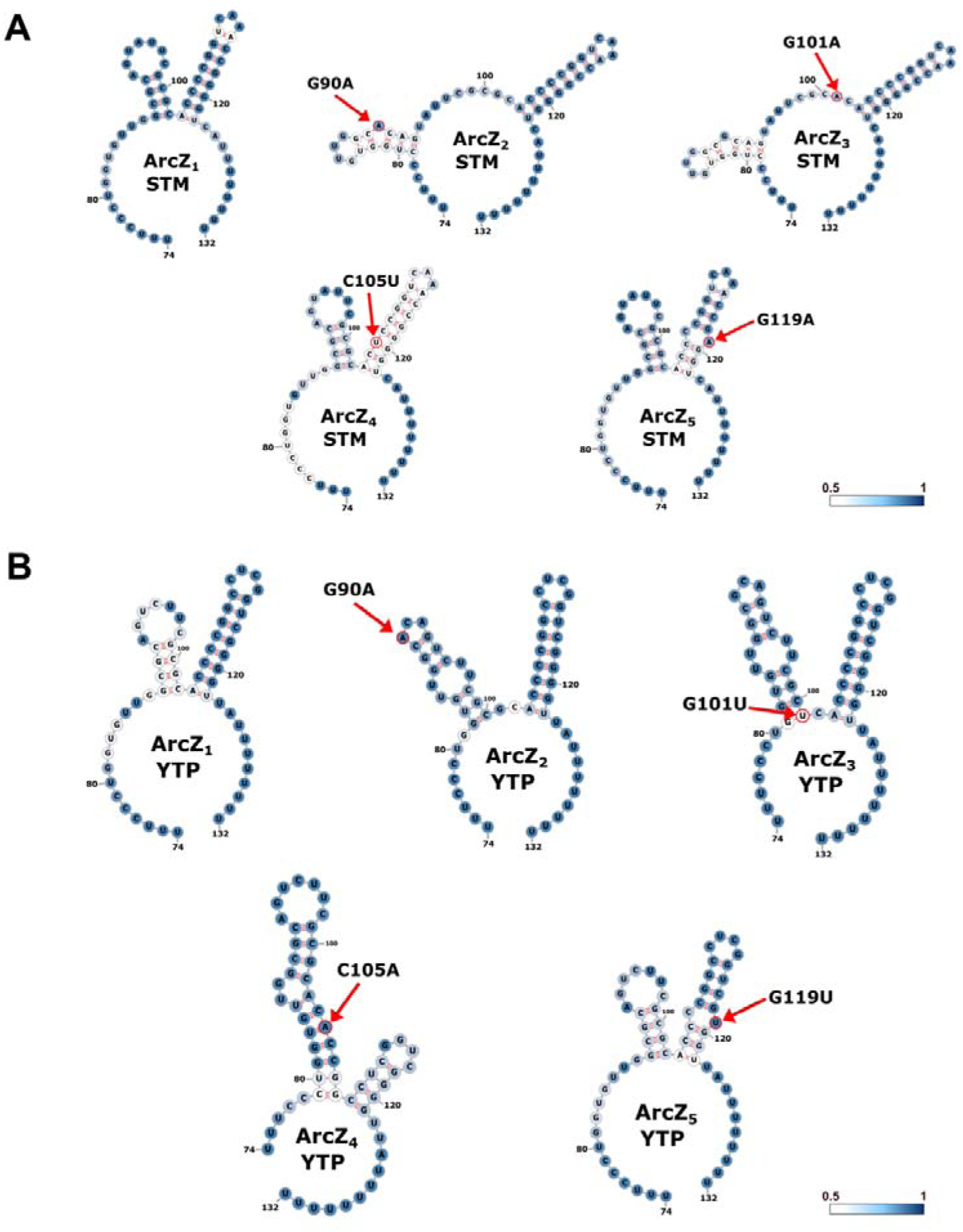
Predicted secondary structures of ArcZ variants in *S. enterica* and *Yersinia sp*. Structures were predicted using RNAfold v2.0 (ViennaRNA Package 2.0). The blue gradient indicates base-pairing probability (0.5-1.0); red arrows indicate substitution positions. For each species, the wild-type structure (ArcZ_1_) is shown alongside structures carrying substitutions at positions equivalent to those identified in *D. solani*. **(A)** Predicted structures of the processed ArcZ sRNA (59 nt) from *S. enterica* LT2 wild-type (ArcZ_1_ STM) and variants carrying substitutions homologous to those identified in *D. solani* (G90A, G101A, C105T, G119A). Note that position 101 carries a G➔A substitution in *S. enterica*, reflecting the naturally occurring variant at this position in this species (Fig. S7B). **(B)** Predicted structures of the processed ArcZ sRNA (59 nt) from *Yersinia sp.* NCTC 10275 wild-type (ArcZ_1_ YTP) and homologous variants (G90A, G101T, C105T, G119T). Substitutions at positions 101 and 119 are G➔T in *Yersinia sp.* but G➔A in *S. enterica*, consistent with the natural polymorphisms observed at these positions across *Enterobacterales* isolates (Fig. S7B).

To dissect this pathway further, we deleted genes involved in its individual enzymatic steps. The acetoin/2,3-butanediol pathway converts two pyruvate molecules into 2,3-butanediol through three sequential reactions: BudB condenses two pyruvate molecules into acetolactate, BudA converts acetolactate into acetoin with CO_2_ release, and BudC reduces acetoin to 2,3-butanediol using NADH as a cofactor (Fig. 5C). Deletion of *budAB*, which prevents acetoin formation, conferred a significant competitive advantage against the *arcZ_1_* strain, whereas deletion of *budC*, which blocks only the final conversion of acetoin into 2,3-butanediol, did not (Fig. 5B). This genetic dissection points to BudAB-dependent acetoin pathway activity, rather than 2,3-butanediol production alone, as a cooperative function exploited by *arcZ* variants cheaters.

Consistent with this interpretation, acetoin production, measured in LB supplemented with 0.5% glucose, a condition selected to stimulate fermentative metabolism and maximize BudAB-dependent output under controlled *in vitro* conditions, was significantly reduced in all five *arcZ* variants relative to that of the *arcZ_1_* strain (Fig. 5D). Moreover, the competitive advantage of the Δ*arcZ_1_*strain was reduced approximately by half when competed against Δ*budR* or Δ*budAB* strains instead of the *arcZ_1_* strain, although it remained positive (Fig. 5B). This reduction was accompanied by fewer generations completed by the Δ*arcZ_1_*strain in the presence of Δ*budR* or Δ*budAB* strains than in the presence of the *arcZ_1_* strain (Fig. S6D). These results indicate that BudAB-dependent pathway activity in the surrounding population is a major, but not exclusive, contributor to the Δ*arcZ_1_* strain cheating advantage.

The functional importance of this pathway was further supported by maceration and pH assays. The Δ*arcZ_1_*, Δ*budR*, and Δ*budAB* strains all displayed strongly reduced tuber maceration in monoculture, whereas the Δ*budC* strain retained near-wild-type maceration (Fig. 5E). Similarly, infected tissue pH remained higher in *arcZ_1_*and Δ*budC* strains than in Δ*arcZ_1_*, Δ*budR*, and Δ*budAB* strains during infection (Fig. 5F), indicating that loss of ArcZ function phenocopies disruption of the BudAB-dependent branch of the pathway. Thus, BudAB-dependent activity is associated both with virulence in monoculture and with maintenance of a less acidic infection environment.

Supporting this interpretation, quantitative proteomics of the Δ*arcZ_1_* strain revealed increased abundance of proteins associated with alternative pyruvate-derived fermentation routes, including FocA, D-lactate dehydrogenase, and GrcA (Fig. S5B, Table S4). Consistently, *grcA* was also upregulated in the transcriptomic datasets of all *arcZ* variants except *arcZpmut* (Table S3). These changes are consistent with a metabolic shift away from BudAB-dependent acetoin production and toward acidic fermentation routes in ArcZ-deficient cells, although the corresponding metabolites remain to be directly quantified.

Together, these results identify BudAB-dependent acetoin pathway activity as a major cooperative function exploited by *arcZ* variants cheaters during potato tuber infection. ArcZ-proficient cells contribute to acetoin production, tuber maceration, and maintenance of a less acidic infection environment, whereas ArcZ-deficient variants reduce their contribution to this pathway while retaining a frequency-dependent benefit in mixed populations.

## Discussion

In contrast with bacterial social cheating strategies that arise through mutations in protein-coding genes such as transcriptional regulators (7, 24), we show that cheating during host infection can emerge through mutational modulation of the conserved post-transcriptional regulator, ArcZ. Point mutations in *arcZ* satisfy the four canonical criteria for social cheating. All five variants gain a fitness advantage exclusively during co-infection with *arcZ_1_* cells, with no benefit detectable *in vitro*. This advantage is maximal when variants are rare and declines as their frequency increases, fulfilling the criterion of negative frequency-dependence (1, 2). Total bacterial productivity declines with increasing cheater frequency, consistent with a tragedy of the commons (4, 25). Finally, the competitive advantage of the non-functional *arcZ_2_* variant is significantly lower against the partial-loss-of-function *arcZ_3_* cooperator than against *arcZ_1_* strain which produce a functional ArcZ. This establishes a graded relationship between cooperator output and cheater benefit. Together, these properties make alternative explanations for negative frequency-dependence unlikely, since the graded reduction of cheater advantage against a partial-loss-of-function cooperator is a diagnostic signature of public good exploitation that niche partitioning, cross-feeding, density-dependent effects, or spatial dynamics cannot account for (26–28).

Having established that *arcZ* variants behave as social cheaters, we next identified the cooperative function they exploit. Transcriptomic profiling identifies a core of 186 genes consistently downregulated across all five *arcZ* variants, enriched in secreted and diffusible functions. Systematic genetic dissection establishes that BudAB-dependent acetoin production is the dominant cooperative function exploited during infection: deletion of the solanimycin biosynthetic cluster, the chrysobactin cluster, or the combined pectate lyase and metalloprotease double mutant conferred no competitive advantage, whereas deletion of *budAB* or *budR* did. The mechanism is linked to tissue pH: *arcZ_1_* cells divert pyruvate flux toward acetoin and 2,3-butanediol via the BudAB pathway, limiting acidic fermentation end-product accumulation and maintaining a less acidic infection environment (14). Variants that repress BudAB are predicted to redirect pyruvate toward mixed-acid fermentation end-products. Consistent with this, the Δ*arcZ_1_*strain proteome shows increased abundance of FocA, D-lactate dehydrogenase, and GrcA, proteins associated with formate transport, D-lactate production, and pyruvate formate-lyase activity, suggesting that repression of the Bud pathway may favor mixed-acid fermentation routes, though direct metabolite quantification during infection remains to be performed (29).

This identification of acetoin-mediated pH buffering as an exploitable cooperative public good situates our findings within a broader pattern across bacteria. The same cooperative output finds a parallel in *Vibrio cholerae*, where the cooperative production of acetoin and 2,3-butanediol to maintain environmental pH is regulated by the Qrr sRNAs downstream of quorum sensing (30). Strikingly, the regulatory logic is inverted: whereas ArcZ activates acetoin production in *D. solani* and its loss-of-function generates cheater variants, constitutive overexpression of Qrr sRNAs in *V. cholerae*, which mimics a quorum sensing-deficient state, imposes a cheating behavior on the overexpressing strain by abolishing its contribution to acetoin-mediated pH buffering (30). Despite this regulatory inversion, both systems converge on the same metabolic public good, the diversion of pyruvate flux toward pH-neutral acetoin, underscoring that pH buffering via acetoin production is a recurrently exploitable cooperative output across phylogenetically distinct bacteria regardless of the regulatory architecture that controls it. A related phenomenon has been described in clonal populations of *Bacillus subtilis*, where a subpopulation of cells producing acetoin to neutralize extracellular acetate maintains a less acidic growth environment from which non-producing cells benefit without contributing to pH buffering (31). This extends the susceptibility of acetoin-mediated pH maintenance to exploitation beyond *Enterobacterales* and shows that it is independent of the genetic mechanism that generates phenotypic heterogeneity. These *arcZ* variants thus benefit from pH buffering they do not contribute to, extending the repertoire of exploitable cooperative traits beyond diffusible macromolecules such as siderophores and exoenzymes. The residual competitive advantage of the Δ*arcZ_1_* strain against the Δ*budAB* mutant indicates that additional ArcZ-dependent functions contribute to cheater fitness. The solanimycin cluster, while repressed in all variants, did not contribute detectably in the potato tuber context, but may be more important in soil or polymicrobial environments where intermicrobial competition is stronger.

How individual substitutions produce a graded loss of function is explained by their distinct structural consequences. The two functional classes defined by the *arcZ* allelic series reflect distinct molecular mechanisms of ArcZ impairment. For the complete loss-of-function class (*arcZ_2_*, *arcZ_5_*), G90A and G119T induce pronounced structural perturbations in the central stem and terminal stem-loop respectively, reducing base-pairing probability at these positions and impairing RNase E-mediated processing, as evidenced by accumulation of the unprocessed full-length species and near-complete loss of the processed 59 nt form. Since the processed form is the regulatory-active species required for Hfq-dependent mRNA base-pairing (15, 16), its absence accounts for the complete loss of ArcZ-dependent outputs in these variants. For the partial loss-of-function class (*arcZ_3_*, *arcZ_4_*), G101T and C105T yield predicted secondary structures closely resembling ArcZ_1_, yet retain only partial processed ArcZ accumulation and display intermediate phenotypes. The mechanism by which these structurally conservative substitutions impair ArcZ function remains unresolved: they may reduce the efficiency of RNase E recognition at the processing site, alter Hfq binding affinity through subtle structural changes in the processed sRNA not captured by thermodynamic modeling, or modify seed-region base-pairing with specific target mRNAs, thereby altering target regulation without affecting overall folding. Distinguishing between these possibilities would require systematic *in vivo* RNA interactome analysis and structural probing of the processed sRNA variants in the presence and absence of Hfq.

Beyond their structural consequences, the recurrence of identical substitutions across phylogenetically distant genera indicates that these positions are recurrently targeted during evolution. The four positions mutated in *D. solani* are naturally polymorphic in *S. enterica* and *Yersinia* isolates, where equivalent substitutions occur at the same residues (Fig. S7). The same sRNA region is naturally polymorphic across several *Dickeya* species (Fig. S8), indicating that it tolerates variation both within the genus and across more distant *Enterobacterales*. As shown here in *D. solani*, substitutions at G90 and G119 severely disrupt the processed ArcZ fold and abolish regulatory function, whereas substitutions at G101 and C105 preserve partial structural integrity and yield intermediate phenotypes. Secondary structure predictions for equivalent substitutions identified in naturally occurring *S. enterica* and *Yersinia* isolates suggest similar structural perturbations (Fig. S9), but whether these variants impair ArcZ processing and function in these species to the same extent as in *D. solani* has not been tested experimentally.

This evolutionary recurrence intersects with an established role for ArcZ in pathogen virulence. In plant pathogens of the Pectobacteriaceae family, loss of ArcZ function has been shown to reduce virulence. In *D. dadantii*, ArcZ directly represses *pecT*, thereby derepressing the RsmB-RsmA regulatory cascade and promoting production of the T3SS and pectate lyases; and an *arcZ* deletion strain displays drastically reduced virulence (21). Consistently, *pecT*, encoding a LysR-type repressor of PCWDE production, is upregulated in some *arcZ* variants in our transcriptomic datasets. In *D. dadantii*, ArcZ directly base-pairs with the *pecT* 5’ UTR to relieve PCWDE repression, requiring the processed ArcZ form (21). Whether an equivalent interaction occurs in *D. solani* remains to be tested. However, its functional consequences for PCWDE production and virulence in this species are unclear, as pectinase and cellulase activities were not significantly reduced in the tested variants (Fig. S2D, S2E). In other pathogens, a *Pectobacterium carotovorum* mutant lacking *arcZ* is similarly attenuated, although its molecular targets remain uncharacterized (32). In *Erwinia amylovora*, ArcZ is required for full virulence, partly through modulation of resistance to plant-derived oxidative stress (33, 34). In *Salmonella* Typhimurium, ArcZ acts as a pleiotropic post-transcriptional regulator: it represses targets including *sdaC*, tpx, the chemotaxis determinant STM3216 (16), and also *hilD*, the master activator of the SPI1-encoded T3SS, in an oxygen-dependent manner that links environmental sensing to the control of virulence gene expression during host colonization (35). However, the direct contribution of ArcZ loss-of-function to reduced virulence *in vivo* has not been established in this species. In *Salmonella*, where ArcZ restricts T3SS expression (35) and where T3SS-deficient variants are established social cheaters during co-infection (36), the functional logic would be directly analogous. Despite this broad association between ArcZ function and virulence-associated outputs, no study had previously documented a role for natural *arcZ* mutations in generating competitive variants during host infection. The present study directly tests and confirms this prediction in a plant pathogen.

The pleiotropic cost-imposing role of ArcZ across *Enterobacterales*, combined with the mutational accessibility of the positions we identify here, raises the possibility that social cheating through single-nucleotide mutations in a regulatory sRNA is not restricted to plant pathogens. Systematically investigating *arcZ* allelic variation in animal pathogens such as *Salmonella* and *Yersinia* during host infection now appears as a compelling next step to determine whether this mutational route to social cheating operates broadly across Enterobacterales. More immediately, the recurrent and independent emergence of distinct *arcZ* alleles in all three experimental evolution replicates, combined with the absence of variants *in vitro*, supports host-specific selective pressure at this locus. Pleiotropic sRNAs, by virtue of their post-transcriptional control over multiple costly traits through a single compact molecule, represent a previously unrecognized class of mutational targets for the evolution of social cheating in bacterial pathogens.

## Materials and Methods

A full list of bacterial strains, plasmids, and oligonucleotides used in this study is provided in SI Appendix, Tables S1, S5 and S6. Detailed descriptions of all experimental procedures and associated references are provided in SI Appendix, Materials and Methods. These include bacterial growth conditions, strain construction and genetic engineering, yeast growth inhibition assays, extracellular enzyme activity assays, Northern blot analysis, potato tuber virulence and pH measurements, transcriptional reporter assays, competition experiments, in vivo and in vitro experimental evolution, amplicon sequencing and variant frequency analysis, whole-genome sequencing, RNA-seq, quantitative proteomics, acetoin quantification, RNA secondary structure prediction, sequence alignment and homology searches, and statistical analyses. Amplicon sequencing and whole-genome sequencing datasets, together with the custom analysis script, are available through Figshare at https://doi.org/10.6084/m9.figshare.31007098.v1. RNA-seq data have been deposited in the NCBI Gene Expression Omnibus under accession no. GSE334754. Mass spectrometry proteomics data have been deposited in the MassIVE repository under accession no. MSV000102049.

## Acknowledgments and Funding Sources

We thank Jan van der Wolf (Wageningen University) for providing the original IPO 2222 cryotube stock, which was essential for investigating the population heterogeneity within the reference strain. We also thank Sonia Humphris (James Hutton Institute) and Robert Czajkowski (University of Gdansk) for providing the IPO 2222 isolates used for comparative phenotypic and genotypic analyses. We acknowledge the contribution of SFR Biosciences (Université Claude Bernard Lyon 1, CNRS UAR3444, Inserm US8, ENS de Lyon) Protein Science Facility, especially Frédéric Delolme and Adeline Page, for the mass spectrometry analyses. We gratefully acknowledge support from the CNRS/IN2P3 Computing Center (Lyon, France) for providing computing and data-processing resources. This work was supported in part by the Agence Nationale de la Recherche: grant ANR-22-CE35-0017 attributed to E.G., grants ANR-17-CE11-0009 and ANR-25-CE11-6139 attributed to L.A. K.P. acknowledges funding by the DFG (EXC 2051 – Project number 390713860) and the European Research Council (ArtRNA, CoG-101088027).

## Author contributions

Q.D., L.A. and E.G. designed research; Q.D., T.B., M.S. and V.U. performed research; Q.D., T.B., M.S., R.M., J.C., K.P. L.A and E.G. analyzed data; A.R., K.P., L.A. and E.G. provided funding and Q.D., L.A. and E.G. wrote the paper, with contribution from M.S. and K.P., and all other authors.

## Tables S1-S7 legends

**Table S1. *Dickeya solani* strains and genomic variants identified in this study.**

This table details the origin of each sequenced strain and the reference genome used for variant calling (IPO 2222, GenBank accession no. NZ_CP015137.1). Each row corresponds to one sequencing experiment (EW identifier). For each detected variant, the table provides its genomic coordinate, the affected gene or genomic region, the nucleotide substitution, and the predicted protein-level consequence, where applicable. Variants at the *arcZ* locus are classified as *arcZ_1_*, *arcZ_2_*, *arcZ_3_*, *arcZ_4_*, *arcZ_5_* and *arcZ_pmut_* and were used to assign strains to distinct IPO 2222 sub-lineages. *arcZ_1_*is the allele of reference strain DS624. Ø mean no mutation detected or no predicted protein-level consequence.

**Table S2. Illumina amplicon sequencing analysis of the *arcZ* locus following experimental evolution.**This table presents deep-sequencing data from PCR-amplified *arcZ* fragments obtained from the original IPO 2222 reference cryotube from the Netherlands collection, the initial inoculum, and bacterial populations passaged for approximately 30 generations in LB medium or through two successive potato tuber infections. The experiment included four biological replicates for the LB condition and three biological replicates for the potato tuber condition, with each biological replicate analyzed using two technical PCR replicates. For each sample, the table reports the total number of reads mapped to the *arcZ* locus and the frequency of the wild-type *arcZ_1_* allele. The two most abundant mutant sequences, designated Top read 1 and Top read 2, are described by their nucleotide variation, absolute read count, and percentage of total mapped reads.

**Table S2. Illumina amplicon sequencing analysis of the *arcZ* locus following experimental evolution.** Deep-sequencing data are reported for four sample types: inoculum, populations passaged for ∼30 generations in LB medium (four biological replicates; LB1–LB4), populations recovered after two successive potato tuber infection cycles (three biological replicates; PDT1–PDT3), and the original IPO 2222 reference cryotube (Netherlands Plant Disease Collection). Each biological replicate was analyzed in two independent PCR replicates (denoted -1 and -2). For inoculum, LB and PDT samples, columns report the read count and percentage of total mapped reads for the *arcZ_1_* allele, the nucleotide change and read count of the two most abundant mutant sequences (Top 1 and Top 2), and total mapped reads; percentages are given in parentheses. *Other* indicates reads not assignable to *arcZ_1_* or the two dominant mutants. For cryotube samples, named allele columns (*arcZ_1_*, *arcZ_2_*, *arcZ_3_*) replace the Top 1/Top 2 scheme, and frequencies are expressed as decimal fractions. The double promoter substitution at positions −60/−59 is noted as CT➔ AA (corresponding to the arcZ*_pmut_* allele). Trimmed FASTQ datasets were analyzed using a custom Python script to identify deletions, substitutions, and insertions in *arcZ* sequence.

**Table S3. RNA-seq transcriptomic analysis of the five *arcZ* variants relative to the *arcZ_1_* reference strain.**

This table presents differential gene expression data for the *arcZ_2_*, *arcZ_3_*, *arcZ_4_*, *arcZ_5_* and *arcZ_pmut_*variants relative to the *arcZ_1_* strain DS624. Strains were grown in LB medium at 30°C to an OD_600_ of 0.5, with three biological replicates analyzed per strain. For each variant, separate worksheets report significantly upregulated and downregulated genes, including locus identifiers, gene names, functional annotations, log_2_ fold changes, and FDR-adjusted P values. The « CORE_DOWN » and « CORE_UP » worksheets list genes that were significantly downregulated or upregulated, respectively, in all five *arcZ* variants compared with *arcZ_1_*. Differentially expressed genes was assessed using the edgeR algorithm with TMM normalization and using a threshold of log_2_ fold change ≤ -1 or ≥ 1 and an FDR-adjusted P value < 0.05.

**Table S4. Quantitative proteomic analysis of the Δ*arcZ_1_* mutant relative to the *arcZ_1_* reference strain.**

Strains DS354 (Δ*arcZ_1_*) and DS49 (*arcZ_1_*) were grown in M63 medium with 1% sucrose at 30°C to OD_600_ = 0.3 (three biological replicates each); proteins were identified and quantified by LC-MS/MS using Proteome Discoverer 2.5. The proteomic_data worksheet contains the full Proteome Discoverer output (UniProt accessions, PEP scores, sequence coverage, peptide counts). The analysis worksheet reports differential abundance results cross-referenced to IPO 2222 locus tags, with log_2_ fold change and FDR-adjusted P value for each protein; significance thresholds were log_2_ fold change, ≤ -1 or ≥ 1 and FDR-adjusted P < 0.05 (one-way ANOVA). A total of 106 proteins showed reduced abundance and 50 showed increased abundance in Δ*arcZ_1_*.

**Table S5. Bacterial strains and plasmids used in this study.**

All *Dickeya solani* and *Escherichia coli* strains and plasmids used in this study are listed with their genotype, relevant genetic modifications, antibiotic-resistance markers, *arcZ* allele (for *D. solani* strains), and source or reference.

**Table S6. Oligonucleotides used in this study.**

Sequences are given in the 5′-to-3′ orientation. Lowercase letters denote genomic target-specific regions; uppercase letters denote adapter or universal sequences.

**Table S7. Bacterial strains and genome accession numbers used for comparative ArcZ sequence analyses.**

This table lists the bacterial species and strains included in the comparative analysis of ArcZ sequences across *Dickeya*, *Salmonella*, *Yersinia*, *Shigella*, and *Serratia*. For each strain, the table provides the strain designation, species or subspecies, sequence database, and genome accession number or database identifier used to retrieve the *arcZ* locus. Sequences presented in Figures S7 and S9 were retrieved from NCBI, whereas those presented in Figure S8 were retrieved from ASAP. These sequences were used for the nucleotide alignments and predicted secondary-structure comparisons shown in Figures S7, S8 and S9.

