## Supplemental Material and Methods for "Host infection selects for sRNA variants that drive bacterial social cheating"

**SI Appendix, Materials and Methodsw**

**Bacterial Strains, Plasmids, and Growth Conditions**

The *Escherichia coli* and *Dickeya solani* strains, plasmids and oligonucleotides used in this study are listed in Table S5 and Table S6. The genome accession number for *D. solani* IPO 2222 is CP015137.1. *E. coli* was routinely grown at 37°C in Luria‑Bertani (LB) broth. *Kluyveromyces lactis* was grown at 30°C in YPG rich medium (1% Bacto yeast extract, 1% Bacto peptone, 2% glucose). *D. solani* strains were cultivated in LB unless otherwise specified. For yeast inhibition assays, *D. solani* strains were grown in M63 minimal medium supplemented with 1% sucrose (2 g (NH_4_)_2_SO_4_, 13.6 g KH_2_PO_4_, 2.5 mg FeSO_4_ 7H_2_O, 0.2 g MgSO_4_ 7H_2_O, 10 g sucrose, per liter). For growth curve measurements, strains were grown in 200 µL M63 supplemented with 0.2% glycerol in 96-well plates placed inside a temperature‑controlled plate reader (Tecan Infinite M200 PRO) with intermittent shaking (100 rpm, 60 s every 15 min) at 30°C, and OD_600_ was monitored every 15 min.

When required, antibiotics were added at the following concentrations: ampicillin (Amp), 100 µg/mL; streptomycin (Sm), 20 µg/mL; gentamicin (Gm), 10 µg/mL and chloramphenicol (Cm), 4 µg/mL; for both *D. solani* and *E. coli*. Diaminopimelic acid (DAP) was supplemented at 57 µg/mL for growth of the *E. coli* MFD*pir* strain. Solid media contained 12 g/L agar.

**Construction of Gentamicin-marked *D. solani* Strains**

For competition experiments, all tested strains were marked with a gentamicin resistance cassette by inserting a mini‑Tn7‑Gm cassette into the attTn7 site using the pTn7‑M and pTnS3 plasmids, as previously described for *D. solani* D s0432‑1 (1). Correct integration of the Gm^R^ cassette was verified by colony PCR using the primer pair L365/L848 (yielding a 350 bp amplicon).

**In‑Frame Deletion Mutants**

In‑frame deletion mutants were constructed using the *sacB* counter‑selection method (2). To facilitate the selection of recombinant plasmids during cloning, the suicide vector pRE112‑*lacZ*α was constructed to enable blue/white screening. The parental vector pRE112, an R6K‑based suicide plasmid carrying the *sacB* and *cat* genes, was linearized with SacI and KpnI. A 599‑bp fragment containing the *lacZ*𝛼 open reading frame and its upstream regulatory elements was PCR‑amplified from pGNW6 (3) using primers L1982 and L1983. The linearized backbone and the *lacZ*𝛼 cassette were then joined via TLTC (4) to replace the original SacI‑KpnI polylinker.‬‬‬‬‬‬‬‬‬

To make the recombinant suicide plasmids, two PCR fragments corresponding to the upstream and downstream approximately 500 bp flanking regions of the target gene or cluster were cloned into pRE112 or pRE112‑lacZ𝛼, linearized using the restriction enzymes SacI/KpnI or primers L758/L1227 respectively, using TLTC or Gibson assembly. Chemically competent *E. coli* DH5α λ*pir* cells were transformed using the Mix and Go kit (Zymo Research) and transformants were selected on LB agar supplemented with chloramphenicol. Plasmids were verified by colony PCR with primers L762/L763, restriction digestion, and Sanger sequencing (Eurofins). Plasmids were then transferred into *E. coli* MFD*pir* (5) by transformation and subsequently introduced into *D. solani* by conjugation. For conjugation, equal volumes of *D. solani* and *E. coli* MFD*pir* cultures were mixed, centrifuged, resuspended in 90 µL LB supplemented with DAP, and spotted onto LB agar incubated at 30°C for 18 h. Bacteria were then resuspended in 1 mL LB, serially diluted, and plated on LB agar supplemented with chloramphenicol to select first-recombination events. Transconjugants were subsequently plated on LB agar without NaCl supplemented with 5% sucrose and incubated at 19°C for 2 to 3 days to select second-recombination events. Sucrose‑resistant colonies were patched on LB‑Cm plates to verify plasmid loss. All deletions were confirmed by colony PCR using flanking primers listed in Table S6.

The following in‑frame deletions were generated. For the solanimycin biosynthetic cluster, genes *solA* through *solL* were deleted as a single continuous fragment. For the chrysobactin siderophore cluster, genes *cbsF*, *cbsH*, *fct*, *cbsC*, *cbsE*, *cbsA*, and *cbsP* were deleted as a single continuous fragment. For the pectate lyase cluster, genes *pelB*, *pelC*, and *pelZ* were deleted. For the metalloprotease cluster, genes *prtB*, *prtC*, and *prtA* were deleted. A Δ*pelBCZ* Δ*prtBCA* double mutant was constructed by applying the deletion procedure of *pelBCZ* to the Δ*prtBCA* single mutant. For the acetoin/2,3‑butanediol pathway, independent deletion mutants of *budR*, *budAB*, and *budC* were generated. Gm^R^‑marked derivatives were constructed from these strains as described above (Table S5).

**Yeast Growth Inhibition Assay**

*D. solani* *arcZ_1_* strain DS624 and *arcZ* variant strains were grown at 30°C with 120 rpm shaking in M63 minimal medium supplemented with 1% sucrose until they reached OD_600_ = 2.0. YPG agar medium was melted, cooled to approximately 45°C, and mixed with a *K. lactis* overnight culture at 2 OD_600_ units per 30 mL agar to prepare seeded plates. Five microliters of each *D. solani* suspension were spotted onto seeded plates and incubated at 30°C for 24 to 48 h prior to visualization of inhibition zones.

**RNA Isolation and Northern Blot Analysis**

For ArcZ quantification, *D. solani* *arcZ_1_* strain DS624 and *arcZ* variant strains were grown at 30°C with 120 rpm shaking in M63 medium supplemented with 1% sucrose to OD_600_ = 0.8. One‑milliliter aliquots were fixed by mixing with 1 mL cold methanol (4°C) and pelleted by centrifugation. Cell pellets were stored at -80°C prior to extraction. RNA was extracted using the RNAsnap method (6). Pellets were resuspended in 100 µL RNAsnap buffer (18 mM EDTA pH 8.0, 0.025% SDS, 95% formamide) and heated twice at 95°C for 7 min. The lysate was mixed with 650 µL Tri‑Reagent (Zymo Research), and RNA was purified using the Direct‑zol RNA MiniPrep kit (Zymo Research) according to the manufacturer's instructions. RNA was eluted in 30 µL DNase/RNase‑free water and quantified by NanoDrop spectrophotometry.

For Northern blot analysis, 1.2 µg of total RNA was resolved per lane on an 8% acrylamide-urea gel (Invitrogen) with 1 µL Low Range ssRNA Ladder (New England Biolabs) as a size standard. Electrophoresis was performed at 200 V in TBE buffer. RNA was transferred to a nylon membrane (Hybond-N+, GE Healthcare) by electrophoretic transfer in 0.5x TBE (30 min at 300 mA, Transblot, Bio‑Rad) and crosslinked by UV irradiation (254 nm, Stratalinker, Stratagene). Membranes were pre‑hybridized for 4 h at 42°C in ULTRAhyb buffer (Ambion) and hybridized overnight at 42°C with the biotinylated probe LA196 (5'-Biotin-AAAAAAAATGACCCCGACCGAGGTCGGGGTGCGCGAATTATGCGCCAACACCAGGGAAAGCG-TgBiotin-3', purchased from Sigma‑Aldrich). Membranes were washed at 65°C with 2x SSC containing 0.1% SDS. Hybridized probes were detected using the Chemiluminescent Nucleic Acid Detection Module (Pierce) with streptavidin‑HRP and luminol. Chemiluminescence signals were acquired using a CCD camera imaging workstation (Thermo Scientific).

**Virulence Assay on Potato Tuber**

Overnight cultures of *D. solani* *arcZ_1_* strain DS624 and *arcZ* variant strains grown in LB broth were harvested by centrifugation, washed in 10 mM MgSO_4_, and resuspended to 8 x 10^7^ bacteria/mL. Potato tubers were wounded to a depth of approximately 2 cm using a sterile pipette tip and inoculated with twenty-five microliters of bacterial suspension (total inoculum: 2 x 10^6^ bacteria) and sealed with mineral oil. After 48 h of incubation at 30°C, macerated tissue was collected and weighed to quantify virulence. Four biological replicates were performed per strain.

For pH monitoring, a pH probe was inserted directly into the maceration zone of each infected tuber at 0, 24, and 48 h post-infection and pH was recorded. Four biological replicates were performed per strain.

**Construction of transcriptional fusions and measurement of the luminescence**

To monitor *arcZ* promoter activity, the promoter regions of *arcZ_1_* and *arcZ_pmut_* from DS624 and DS945 were amplified by PCR using primers L2580/L2581 and cloned by HiFi assembly into pSEVA421‑*luxCDABE*‑*gfp* previously linearized with primers L2590/L2591. Plasmids pSEVA421‑*promo arcZ*(WT)::*luxCDABE*‑*gfp* and pSEVA421‑*promo arcZ*(CT🡪AA)::*luxCDABE*-*gfp* were verified by restriction digestion and Sanger sequencing (Table S6). For luminescence assays, bacterial cultures were inoculated at OD_600_ = 0.006 in 96‑well plates in M63 medium supplemented with 1% sucrose and incubated at 30°C. Luminescence was measured at OD_600_ = 0.3 using a Tecan Infinite M200 plate reader with an integration time of 1 s per well. Luminescence values were normalized to OD_600_. Three biological replicates were performed.

**Competition Assays**

Overnight cultures of D. solani strains grown in LB broth were harvested by centrifugation, washed in 10 mM MgSO_4_, and resuspended to 8 x 10^7^ bacteria/mL. Competition mixtures were prepared by combining gentamicin‑resistant (Gm^R^) strains with unmarked gentamicin‑sensitive (Gm^S^) competitors at the indicated initial ratios. Unless otherwise specified, Gm^R^ strains were inoculated as the minority population at a 1:100 Gm^R^:Gm^S^ ratio by mixing 8 µL of the Gm^R^ strain with 800 µL of the Gm^S^ competitor.

For competition assays involving arcZ variants, Gm^R^ strains DS899 (arcZ_2_), DS901 (arcZ_3_), DS947 (arcZ_4_), DS948 (arcZ_5_), and DS949 (arcZ_pmut_) were competed against the unmarked arcZ_1_ strain DS624. For competition assays involving the ΔarcZ_1_ deletion mutant, Gm^R^ strain DS937 (ΔarcZ_1_, D s0432‑1 background) and DS943 (ΔarcZ_1_ complemented with pEGL334 carrying arcZ_1_ (12)) were competed against the unmarked arcZ_1_ strain DS49 (D s0432‑1). This pairing ensures that competitive outcomes reflect differences at the arcZ locus exclusively, as DS937 and DS49 are isogenic except at this locus. To assess the effect of initial strain frequency on competitive dynamics, mixtures were prepared at Gm^R^:Gm^S^ ratios of 1:100, 10:100, 100:100, 100:10, and 100:1 by adjusting volumes accordingly.

For potato tuber competition assays, twenty-five microliters of each bacterial mixture (total inoculum: 2 x 10^6^ bacteria) was inoculated into wounded tubers as described above. After 48 h of incubation at 30°C, macerated tissue was collected, weighed, homogenized, and serially diluted in 10 mM MgSO_4_. For *in vitro* competition assays, twenty-five microliters of the same bacterial mixtures was inoculated into 5 mL of M63 minimal medium supplemented with 0.2% glycerol and incubated at 30°C for 24 h.

To monitor competition dynamics over time, co‑infections were performed using an initial 1:100 ΔarcZ_1_:arcZ_1_ ratio between DS937 (∆*arcZ_1_* Gm^R^) and DS49. twenty-five microliters of bacterial mixture (total inoculum: 2 x 10^6^ bacteria) was inoculated into a wounded potato slice. Infected tissue was sampled at 12, 24, 36, and 48 h post-infection by homogenization followed by plating on selective and non‑selective media.

Bacterial populations were quantified by plating serial dilutions on non‑selective LB agar and LB agar supplemented with gentamicin (10 µg/mL) using an easySpiral automatic plater (Interscience). Colonies were enumerated after 48 h at 30°C using a Scan 1200 colony counter (Interscience). Gm^R^ populations were quantified on gentamicin‑containing plates and total populations on non‑selective plates. Gm^S^ populations were calculated by subtracting Gm^R^ CFU from total CFU.

The Competitive Index (CI) was calculated as:

CI = (Gm^R^ final / Gm^S^ final) / (Gm^R^ initial / Gm^S^ initial)

Results are expressed as log_10_ CI. The number of generations completed by each strain was calculated as (log_10_N_f_ - log_10_N_0_) / log_10_2, where N_0_ and N_f_ are the initial and final CFU counts of that strain, respectively.

***In vivo* Experimental evolution assay during potato tuber infection**

Overnight cultures of *D. solani* *arcZ_1_* strain DS624, a sub-isolate recovered from the IPO 2222 reference cryotube from the Netherlands collection, grown in LB broth were harvested by centrifugation, washed in 10 mM MgSO_4_, and resuspended to 8 x 10^7^ bacteria/mL. Twenty-five microliters of this suspension (total inoculum: 2 x 10^6^ bacteria) were inoculated into wounded potato tubers and the inoculation sites were sealed with a drop of mineral oil (Sigma). After 48 h of infection, macerated tissue was collected and diluted 100‑fold in 10 mM MgSO_4_. Twenty-five microliters of this dilution were inoculated into a new wounded tuber and the inoculation sites were sealed with mineral oil. Two consecutive infection cycles were performed across three independent biological replicates. After two infection cycles, macerated tissue from each replicate was homogenized in 10 mL of 10 mM MgSO_4_. The arcZ region was amplified from the bulk population by PCR using primers L2373/L2374 and variant frequencies were determined by amplicon sequencing as described below. In parallel, serial dilutions were plated to obtain isolated colonies. Two hundred individual colonies per replicate were patched onto YPG agar seeded with K. lactis and incubated at 30°C for 24 h. Colonies failing to inhibit yeast growth were selected and their arcZ locus was sequenced by Sanger sequencing using primers L2373/L2374.

***In vitro* Experimental evolution assay**

Overnight cultures of D. solani arcZ_1_ grown in LB broth were adjusted to OD_600_ = 1.0. Fifty microliters were inoculated into 5 mL fresh LB broth (initial OD_600_ approximately 0.01) and incubated at 30°C with 120 rpm shaking. Upon reaching stationary phase (OD_600_ approximately 2.0), cultures were passaged into fresh LB at the same dilution factor. This process was repeated four times, corresponding to approximately 7.6 generations per passage and a cumulative total of approximately 30 generations. After the fourth passage, the arcZ region was amplified by PCR using primers L2373/L2374 and variant frequencies were determined by amplicon sequencing as described below.

**Amplicon sequencing and variant frequency analysis**

To quantify arcZ variant frequencies in bulk bacterial populations recovered from experimental evolution assays and from the Netherland cryotube, two technical replicate PCR amplicons spanning the arcZ coding region and the arcZ promoter-containing region were generated from genomic DNA using primer pair L2373/L2374. Amplicon library preparation and paired‑end sequencing (2 × 250 bp) were performed by Azenta/Genewiz (Leipzig, Germany) using the NEBNext Ultra II DNA Library Prep Kit (New England Biolabs) according to the manufacturer's instructions. Raw BCL files were converted to demultiplexed FASTQ files using bcl2fastq v2.17.

Paired‑end reads were imported into Galaxy Europe (usegalaxy.eu), merged using PEAR (v0.9.6.4) (7), and adaptor sequences were removed using Cutadapt (v5.2) (8) with the corresponding flanking primer sequences as trimming references, retaining only reads spanning the full amplicon. Trimmed reads were analyzed using a custom Python (v3.11) script implemented with Biopython (v1.86). Reads were oriented relative to the arcZ_1_ reference sequence when required. The script identified and counted reads differing from the arcZ_1_ reference sequence at one or more positions. Single‑nucleotide variants were recorded when reads differed from the reference at a single position. For the arcZ_pmut_ promoter allele, defined by two linked substitutions (CT to AA at positions −60 and −59), reads carrying the complete combination of both substitutions were grouped and counted as a single haplotype. Allele frequencies were calculated as the number of reads supporting a given variant or haplotype divided by the total number of retained reads for that sample. Only variants detected at frequencies ≥ 0.5% were retained for downstream analysis, a threshold set above the expected per‑base error rate of high‑quality Illumina reads after filtering. The Python script, together with the raw NGS datasets, are available at <https://doi.org/10.6084/m9.figshare.31007098.v1>.

**Whole-genome sequencing**

Two milliliters of overnight LB cultures were pelleted and genomic DNA was extracted using the Wizard Genomic DNA Purification Kit (Promega). Long-read sequencing was performed using the Rapid Barcoding Kit 96 (SQK-RBK114.96, Oxford Nanopore Technologies) on a PromethION apparatus with a FLO-PRO114M flow cell (Microsynth). Short-read sequencing was performed on a NovaSeq 6000 platform (2 × 150 bp paired-end, Azenta/Genewiz). Nanopore reads were assembled using Flye. Illumina reads were quality‑checked, trimmed, and used to polish Nanopore assemblies. Whole‑genome comparisons and SNP detection against the reference genome NZ_CP015137.1 were performed using MUMmer (dnadiff) on the Galaxy.eu platform. The raw NGS datasets are available at <https://doi.org/10.6084/m9.figshare.31007098.v1>.

**Global Gene Expression Profiling by RNA-Seq**

D. solani strains DS624 (arcZ_1_), DS623 (arcZ_2_), DS625 (arcZ_3_), DS931 (arcZ_4_), DS944 (arcZ_5_), and DS945 (arcZ_pmut_) were cultivated in triplicate in LB medium at 30°C with 120 rpm shaking. LB medium was selected because sufficient RNA integrity for library preparation and sequencing could not be achieved when strains were cultivated in M63 minimal medium supplemented with 1% sucrose. At OD_600_ = 0.5, transcription was stopped by addition of 0.5 volumes of cold 100% methanol. Total RNA was isolated by hot phenol extraction, treated with Turbo DNase (Thermo Fisher Scientific), and RNA integrity was confirmed using a Bioanalyzer (Agilent). Ribosomal RNA was depleted using rRNA-specific biotinylated probes as previously described (9). rRNA-depleted RNA was purified using Agencourt AMPure XP beads (Beckman Coulter) and fragmented for 5 min at 75°C using the NEBNext Magnesium RNA Fragmentation Module (NEB). cDNA libraries were prepared using the NEBNext UltraExpress RNA Library Prep Kit for Illumina (NEB, E3330L) according to the manufacturer's instructions. Library quality was assessed on an Agilent 2100 Bioanalyzer and pooled libraries were sequenced on a NextSeq 1000 system with 50 nt paired‑end sequencing mode. Demultiplexed reads were trimmed and mapped to the D. solani reference genome (NCBI accession CP015137.1) using the RNA-Seq Analysis tool of CLC Genomics Workbench (Qiagen) with standard parameters (statistical analysis: TMM normalization and edgeR algorithm). Differentially expressed genes were defined as those with a log₂ fold change ≥ 1 or ≤ -1 and an FDR‑adjusted p‑value ≤ 0.05. Three biological replicates were performed per strain. The RNA‑seq data have been deposited in the NCBI Gene Expression Omnibus (GEO) repository under accession number GSE334754.

**Quantitative Proteomics**

Quantitative proteomics was performed on the ΔarcZ_1_ deletion mutant DS354 relative to the arcZ_1_ strain DS49 to provide an independent line of evidence for the ArcZ‑dependent expression program at the protein level. The ΔarcZ_1_ deletion mutant was selected because it represents a complete and unambiguous loss of ArcZ function, providing the strongest signal for identifying ArcZ-dependent proteins independently of the structural effects of individual point mutations present in the different variants. M63 minimal medium supplemented with 1% sucrose was used because it supports robust protein extraction and reproducible quantification under defined nutritional conditions, and because RNA‑seq analysis under these conditions was not feasible due to insufficient RNA quality for library preparation.

DS49 (*arcZ_1_*) and DS354 (Δ*arcZ_1_*) were cultivated in triplicate in M63 minimal medium supplemented with 1% sucrose at 30°C with 120 rpm shaking to OD_600_ = 0.3. Cells were harvested by centrifugation and protein extracts were prepared using the EasyPep™ mini sample preparation kit (Thermo Scientific) following the manufacturer's instructions, including denaturation, reduction, alkylation (10 min, 95°C), and digestion with a LysC/Trypsin enzyme combination (3 h, 37°C), followed by peptide desalting. Peptide concentrations were determined after resuspension in 50 mM TEAB using a quantitative fluorometric peptide assay and samples were labeled with TMT 6‑plex isobaric reagents (Thermo Scientific) according to the manufacturer's protocol. The labeling reaction was quenched with 5% hydroxylamine and labeled peptides were pooled into a single TMT 6‑plex set and desalted on a C18 microcolumn.

After dilution in 0.1% formic acid, the pooled sample was analyzed in three technical replicates on an Ultimate 3000 nanoRSLC system coupled online to a Q Exactive HF mass spectrometer via a nano‑electrospray ionization source (Thermo Scientific). Samples were loaded onto a C18 trap column and separated on a 50 cm C18 analytical column using a 100 min linear gradient from 3.2% to 90% acetonitrile in 0.1% formic acid at 300 nL/min and 40°C. Data were acquired in Data‑Dependent Acquisition (DDA) TOP15 HCD mode, with MS survey scans at a resolution of 120,000 and MS2 fragmentation at a resolution of 45,000 (normalized collision energy = 33). Dynamic exclusion was set to 30 s to limit redundant fragmentation.

Data files were processed with Proteome Discoverer 2.5 (Thermo Scientific) using the SEQUEST HT search engine against the *D. solani* D s0432‑1 proteome (UniProt, June 2023, 5,451 sequences) supplemented with a common contaminants database (116 sequences). Precursor and fragment mass tolerances were set at 10 ppm and 0.02 Da, respectively, with up to two missed cleavages allowed. Oxidation (M), acetylation (protein N-terminus), and TMT6plex (K, peptide N-terminus) were set as variable modifications, and carbamidomethylation (C) as a fixed modification. Peptide and protein identifications were validated using Percolator at a 1% false discovery rate. Relative protein quantification was based on TMT reporter ion abundances; DS354/DS49 abundance ratios were averaged across three technical replicates and statistically validated by ANOVA. Proteins with a log_2_ abundance ratio ≥ 1 or ≤ ‑1 and a FDR P‑value < 0.05 were considered differentially abundant. A total of 2,127 proteins were identified, of which 106 were significantly less abundant and 50 were significantly more abundant in the Δ*arcZ_1_* strain relative to the *arcZ_1_* strain. Proteomics analyses were performed at the Protein Science Facility of SFR Biosciences (UAR3444/CNRS, US8/Inserm, ENS de Lyon, UCBL), Lyon, France. The mass spectrometry proteomics data have been deposited to the Center for Computational Mass Spectrometry repository (University of California, San Diego) via the MassIVE tool with the dataset identifier MassIVE MSV000102049.

**Acetoin quantification assay**

Acetoin production was measured in culture supernatants of DS624 (arcZ_1_), DS623 (arcZ_2_), DS625 (arcZ_3_), DS931 (arcZ_4_), DS944 (arcZ_5_), and DS945 (arcZ_pmut_) as a proxy for BudAB pathway activity. LB medium supplemented with 0.5% glucose was selected to stimulate fermentative metabolism and maximize acetoin production under controlled in vitro conditions. Strains were grown at 30°C with 120 rpm shaking for 16 h. Cultures were centrifuged at 12,000 x g for 2 min and supernatants were collected. Acetoin concentration was determined by the Voges‑Proskauer colorimetric assay (10). Supernatants were diluted 20‑fold in LB to a final volume of 1 mL. Creatine (60 µL, 0.5%), alpha‑naphthol (90 µL, 5% in ethanol), and KOH (60 µL, 40%) were added sequentially, and samples were incubated for 30 min at room temperature with occasional mixing. Absorbance was measured at 515 nm and acetoin concentrations were determined against a standard curve ranging from 0 to 300 µM. Values were normalized to OD_600_ of the corresponding culture. Four biological replicates were performed per strain.

**Secondary structure prediction of ArcZ sRNA variants**

The secondary structures of ArcZ₁ and the four sequence variants ArcZ_2_, ArcZ_3_, ArcZ_4_, and ArcZ_5_ were predicted using RNAfold version 2 (ViennaRNA Package 2.0 ; <https://rna.tbi.univie.ac.at>) (11). The complete sequence of the processed ArcZ sRNA from *D. solani* *arcZ₁* was used as reference input. For each variant, the corresponding single-nucleotide substitution was introduced manually into the reference sequence prior to structure prediction. Because *arcZpmut* carries a promoter mutation leaving the sRNA sequence unchanged, its processed form is structurally identical to ArcZ_1_ and was not modeled separately. Analogous predictions were performed for *S. enterica* LT2 and *Y. pseudotuberculosis* NCTC 10275 ArcZ orthologs with the corresponding substitutions introduced at equivalent positions.

**Nucleotide sequence alignment and homology search**

To determine whether *arcZ* alleles identified in *D. solani* occur naturally in related species, comparative sequence analysis was performed across the *Dickeya* genus and against representative *Enterobacterales*. Wild-type ArcZ sRNA sequences were retrieved from the NCBI nucleotide database for ten *Dickeya* species and for *S. enterica* and *Yersinia* spp. The substitutions identified in *D. solani* *arcZ* variants (G90A, G101T, C105T, G119T) were individually introduced into the corresponding wild‑type sequences and used as queries for BLASTn searches against the NCBI nucleotide database (https://blast.ncbi.nlm.nih.gov) with default parameters. Multiple sequence alignments were performed using ClustalW (12) via the DDBJ ClustalW web server (https://www.genome.jp/tools-bin/clustalw). Alignment outputs were visualized and annotated using Jalview version 2.11.5.1 (13).

**Extracellular Enzyme Activity Assays**

Protease, cellulase, and pectinase activities were assessed on indicator plates. For protease activity, LB agar was supplemented with 0.625% skim milk. For cellulase activity, M63 agar was supplemented with 0.2% carboxymethylcellulose (CMC), 0.08% MgSO_4_, and 0.4% glycerol. For pectinase activity, M63 agar was supplemented with 0.2% glycerol and 0.2% polygalacturonic acid (Dipecta). *D. solani* *arcZ_1_* strain DS624 and *arcZ* variant strains were grown overnight in LB at 30°C and adjusted to OD_600_ = 1.0. Five-microliter drops were spotted onto indicator plates and incubated at 30°C for 24 to 48 h. Pectinase activity was revealed by addition of 10% copper acetate solution onto plate surfaces. Cellulase activity was revealed by staining with 1% Congo Red followed by 1 M NaCl wash. Degradation halo diameters were measured for four biological replicates per strain.

**Statistical Analysis**

All statistical analyses were performed using GraphPad Prism version 10. Comparisons between two groups were performed using the Mann-Whitney test. For growth curves, doubling times were calculated during the exponential growth phase and compared between strains using pairwise Mann-Whitney tests. For proteomic data, differential protein abundance was assessed by ANOVA with a FDR-adjusted p-value threshold of 0.05, as described above. For transcriptomic data, differential gene expression was assessed using the edgeR algorithm with TMM normalization and an FDR-adjusted p‑value threshold of 0.05, as described above.

Results are expressed as mean ± standard deviation (SD) unless otherwise stated. Significance thresholds were set at p < 0.05 (*), p < 0.01 (**), and p < 0.001 (***). Non‑significant differences are indicated as ns.
