## Supplementary material for "Host infection selects for sRNA variants that drive bacterial social cheating": S1 Table

**Table S1**

| **Samples** | **Source of *D. solani* strains** | **Reference  genome** | **Coordinates** | **Gene** | **DNA Mutation** | **Protein modification** | ***arcZ* allele** |
| --- | --- | --- | --- | --- | --- | --- | --- |
| EW1 | DS45 (IPO 2222 strain of the Lyon lab strain collection) | IPO 2222 (NZ_CP015137.1) | 1370270 | NhaR | A848T | Q283L | *arcZ_2_* |
| EW3 | DS470 (IPO 2222 strain of the Gdansk lab strain collection | IPO 2222 (NZ_CP015137.1) | 3882724 | A4U42_RS16650 (anaerobic C4-dicarboxylate transporter DcuB) | G1223A | G408D |  |
| EW4 | DS486 (IPO 2222 - LMG25993) | IPO 2222 (NZ_CP015137.1) | No mutation | ø |  | ø |  |
| EW5 | DS566 (IPO 2222 - NCPPB4479) | IPO 2222 (NZ_CP015137.1) | 1370270 | NhaR | A848T | Q283L |  |
| EW6 | DS620 (IPO 2222 (Scotland, James Hutton Institute) | IPO 2222 (NZ_CP015137.1) | 2530076 | *arcZ* | G101T | ø | *arcZ_3_* |
|  |  |  | 2530087 | *arcZ* | A90G | ø |  |
| EW7 | DS623 (IPO 2222 from Jan Van der Wolff’s lab, Sol and Zms negative) | IPO 2222 (NZ_CP015137.1) | No mutation |  |  | ø | *arcZ_2_* |
| EW8 | DS624 (IPO 2222 from Jan Van der Wolff’s lab, Sol and Zms positive) | IPO 2222 (NZ_CP015137.1) | 2530087 | *arcZ* | A90G | ø | *arcZ_1_* |
| EW9 | DS625 (IPO 2222 from Jan Van der Wolff’s lab, Sol and Zms positive) | IPO 2222 (NZ_CP015137.1) | 2530076 | *arcZ* | G101T | ø | *arcZ_3_* |
|  |  |  | 2530087 | *arcZ* | A90G | ø |  |
| EW10 | DS931 (*arcZ_4_* from DS624 after 2 successive infection in potato tuber) | IPO 2222 (NZ_CP015137.1) | 2530071 | *arcZ* | C105T | ø | *arcZ_4_* |
|  |  |  | 2530087 | *arcZ* | A90G | ø |  |
| EW11 | DS944(*arcZ_5_* from DS624 after 2 successive infection in potato tuber) | IPO 2222 (NZ_CP015137.1) | 2530057 | *arcZ* | G119T | ø | *arcZ_5_* |
|  |  |  | 2530087 | *arcZ* | A90G | ø |  |
| EW12 | DS945(*arcZ_pmut_* from DS624 after 2 successive infection in potato tuber) | IPO 2222 (NZ_CP015137.1) | 2530235 | *arcZ* promoter | T-->A | ø | *arcZ_pmut_* |
|  |  |  | 2530236 | *arcZ* promoter | C-->A | ø |  |
|  |  |  | 2530087 | *arcZ* | A90G | ø |  |
|  |  |  | 3457677 | A4U42_14900 (sugar ABC transporter ATP-binding protein) | C-->T | GUG-->GUU = Valine |  |

Note: Mutations are reported relative to the deposited reference genome NZ_CP015137.1, which itself carries the *arcZ_2_* allele. The functional *arcZ_1_* strains therefore appear as A90G relative to this reference.
