## Supplementary material for "Host infection selects for sRNA variants that drive bacterial social cheating": S5 Table

**Table S5: Bacterial strains and plasmids**

| Bacterial strain and plasmid | Description | Source |
| --- | --- | --- |
| **Strains** |  |  |
| *Escherichia coli* K12 |  |  |
| DH5α | *supE44 lacU169 (*Φ*80lacZ*∆ M15) *hsdR17 (rK mK ) recA1 endA1 gyrA96 thi-1 relA1* | Laboratory collection |
| DH5α λpir | λpir phage lysogen of DH5α | Laboratory collection |
| MFD*pir* | *RP4-2-Tc::(∆Mu1::aac(3)IV-∆aphA-·∆nic35-∆Mu2::zeo) ∆dapA::erm-pir) ∆recA* | (1) |
| *Dickeya solani* |  |  |
| DS49 | *D. solani* D s0432-1 WT *arcZ_1_* | (2) |
| DS354 | DS49 *D. solani* D s0432-1 ∆*arcZ_1_* | (2) |
| DS433 | DS49 *D. solani* D s0432-1 pEGL332 | (2) |
| DS439 | DS354 *D. solani* D s0432-1 ∆*arcZ_1_ /*pEGL332 | (2) |
| DS441 | DS354 *D. solani* D s0432-1 ∆*arcZ_1_ /*pEGL334 | (2) |
| DS45 | *D. solani* IPO2222 WT *arcZ_2_* | (2) |
| DS470 | *D. solani* IPO2222 WT *arcZ_2_* | Strain Robert Czajkowski collection (Gdansk, poland) |
| DS486 | *D. solani* IPO2222 WT *arcZ_2_* | Strain LMG25993 of the BCCM collection |
| DS566 | *D. solani* NCPPB4479 (Biovar 3, Netherlands) *arcZ_2_* | Strain NCPPB collection |
| DS620 | *D. solani* IPO 2222 Scotland *arcZ_3_* | Scotland |
| DS623 | *D. solani* *arcZ_2_* strain from Netherlands cryotube | This study |
| DS624 | *D. solani* *arcZ_1_* strain from Netherlands cryotube | This study |
| DS625 | *D. solani* *arcZ_3_* strain from Netherlands cryotube | This study |
| DS931 | *D. solani* *arcZ_4_* strain from Experimental evolution in potato tuber | This study |
| DS944 | *D. solani* *arcZ_5_* strain from Experimental evolution in potato tuber | This study |
| DS945 | *D. solani* *arcZ_pmut_* strain from Experimental evolution in potato tuber | This study |
| DS936 | DS49 *glmS::Tn7-gent*, Gm^R^ | This study |
| DS937 | DS354 *glmS::Tn7-gent*, Gm^R^ | This study |
| DS941 | DS433 *glmS::Tn7-gent*, Gm^R^ | This study |
| DS942 | DS439 *glmS::Tn7-gent*, Gm^R^ | This study |
| DS943 | DS441 *glmS::Tn7-gent*, Gm^R^ | This study |
| DS947 | DS931 *glmS::Tn7-gent*, Gm^R^ | This study |
| DS948 | DS944 *glmS::Tn7-gent*, Gm^R^ | This study |
| DS949 | DS945 *glmS::Tn7-gent*, Gm^R^ | This study |
| DS899 | DS623 *glmS::Tn7-gent*, Gm^R^ | This study |
| DS900 | DS624 *glmS::Tn7-gent*, Gm^R^ | This study |
| DS901 | DS625 *glmS::Tn7-gent*, Gm^R^ | This study |
| DS999 | DS624 pEGL578 | This study |
| DS1073 | DS624 pEGL579 | This study |
| DS708 | ∆*budAB* | This study |
| DS883 | ∆*budR* | This study |
| DS885 | ∆*budC* | This study |
| DS1020 | *∆budAB glmS::Tn7-gent,* Gm^R^ | This study |
| DS1003 | *∆budR* glmS::Tn7-gent, Gm^R^ | This study |
| DS1004 | ∆budC glmS::Tn7-gent, Gm^R^ | This study |
| DS1029 | ∆*cbsF* | This study |
| DS1030 | ∆*cbsFHCEAP-fct* | This study |
| DS1031 | ∆*pelBCZ* | This study |
| DS929 | ∆*prtBCA* | This study |
| DS1032 | ∆*pelBCZ* ∆*prtBCA* | This study |
| DS1057 | ∆*solABCDEFGHIJKL*(named ∆*sol*) | This study |
| DS1039 | ∆*cbsF* glmS::Tn7-gent, Gm^R^ | This study |
| DS1040 | ∆*cbsFHCEAP-fct* glmS::Tn7-gent, Gm^R^ | This study |
| DS1042 | ∆pelBCZ ∆prtBCA glmS::Tn7-gent, Gm^R^ | This study |
| DS1058 | ∆*solABCDEFGHIJKL*(named ∆*sol)* glmS::Tn7-gent, Gm^R^ | This study |
| **Plasmids** |  |  |
| pTn7-M | Km^R^ Gm^R^, *ori R6K*, *Tn7L* and *Tn7R* extremities, standard multiple cloning site, *oriT* RP4 | (3) |
| pTNS3 | Amp^R^, *ori R6K*, *TnsABCD* operon, *oriT* RP4 | (4) |
| pEGL332 | pWSK29-oriT, Amp^R^ | (2) |
| pEGL334 | pWSK29-oriT-*arcZ_1_*, Amp^R^ | (2) |
| pRE112 | Suicide vector for allelic exchange, Cm^R^, *sacB*, *oriT* RP4, *ori*R6K | (5) |
| pSEVA421 | Sm^R^, *ori RK2, oriT* | (6) |
| pEGL473 | pRE112-lacZ𝛼 | This study |
| pEGL578 | pSEVA421-*promo arcZ*(WT)::*luxCDBAE*-*gfp* | This study |
| pEGL579 | pSEVA421-*promo arcZ* (mut CT->AA)::*luxCDBAE*-*gfp* | This study |
| pEGL434 | pRE112-∆budAB, Cm^R^ | This study |
| pEGL480 | pRE112-∆budR, Cm^R^ | This study |
| pEGL481 | pRE112-∆budC, Cm^R^ | This study |
| pEGL536 | pRE112-lacZ𝛼-∆*prtBCA*, Cm^R^ | This study |
| pEGL588 | pRE112-lacZ𝛼-∆*pelBCZ*, Cm^R^ | This study |
| pEGL589 | pRE112-lacZ𝛼-∆*cbsF*, Cm^R^ | This study |
| pEGL590 | pRE112-lacZ𝛼-∆*cbsFHCEAP-fct*, Cm^R^ | This study |
| pEGL586 | pRE112-lacZ𝛼-∆*solABCDEFGHIJKL*, Cm^R^ | This study |
