## Supplementary material for "Host infection selects for sRNA variants that drive bacterial social cheating": S6 Table

**Table S6: Oligonucleotides used in this study**

| Oligonucleotide | Sequence (5’-3’) | Use |
| --- | --- | --- |
| **L365** | cacagcataactggactgatttc | Primers to check integration of the miniTn7 into the attTn7 site of *D. solani*. |
| **L848** | atgtggcgctgatcaaaggc |  |
| **L2373** | ACACTCTTTCCCTACACGACGCTCTTCCGATCTctccacatccgctgagtacg | Primers used for high-throughput Illumina amplicon sequencing of the *arcZ* locus |
| **L2374** | GACTGGAGTTCAGACGTGTGCTCTTCCGATCTttgaaacacgactggcagcag |  |
| **L1412** | atgccgggcagtcgatg | Primers used for PCR on colonies to amplify *arcZ* region |
| **L1413** | ttgtaagctgcacagagg |  |
| **L1675** | ctcaccggtcagatgattaatgac | Primer used for Sanger sequencing of the *arcZ* region |
| **L758** | TACCCGGGGATCCTCTAGAG | PCR to amplify the linearized pRE112-lacZ$\alpha$ plasmid |
| **L1227** | cgtgactgggaaaaccc |  |
| **L2590** | ctgcaggcatgcaggagg | PCR to amplify the linearized pSEVA421-luxCDABE-GFP plasmid |
| **L2591** | gagctcgaattcgcgcgg |  |
| **L2580** | CGCGGCCGCGCGAATTCGAGCTCaattgcaagcagagtcaagac | PCR to amplify arcZ promoter region to clone in pSEVA421-luxCDABE-GFP |
| **L2581** | TTTTTCCTCCTGCATGCCTGCAGaccatagcaaatacggcgt |  |
| **L2420** | cgccagggttttcccagtcacgggcagatgagcgttaccg | PCR to amplify fragment up of *prtBCA* to clone in pre112-lacZ$\alpha$ |
| **L2421** | tacacaacgaattgttgcatattacttcttccataaagtctg |  |
| **L2422** | atatgcaacaattcgttgtgtaaccggctg | PCR to amplify fragment down of *prtBCA* to clone in pre112-lacZ$\alpha$ |
| **L2423** | ATCCATGGTCGACAAGCTTCTActacggatttatggagaagccc |  |
| **L2640** | acgccagggttttcccagtcacgaatgaagcggacaatggc | PCR to amplify fragment up of *pelBCZ* to clone in pre112-lacZ$\alpha$ |
| **L2641** | gctatgaagtcagatctggaataacggttcattgaa |  |
| **L2642** | tattccagatctgacttcatagcgtgtgttcc | PCR to amplify fragment down of *pelBCZ* to clone in pre112-lacZ$\alpha$ |
| **L2643** | cgaCTCTAGAGGATCCCCGGGTAtacgccttcggccaaaa |  |
| **L2644** | cgccagggttttcccagtcacgtttgtgctggcgcggcgggttg | PCR to amplify fragment up of *cbsF* to clone in pre112-lacZ$\alpha$ |
| **L2645** | tgttaccgacgatgtatcagcgttacctcattgg |  |
| **L2646** | cgctgatacatcgtcggtaacagggataatggc | PCR to amplify fragment down of *cbsF* to clone in pre112-lacZ$\alpha$ |
| **L2647** | gaCTCTAGAGGATCCCCGGGTAgagtccgccctacgtttctc |  |
| **L2648** | cgccagggttttcccagtcacggccgctcaatgagctgaacag | PCR to amplify fragment up of *cbsFHCEAP-fct* to clone in pre112-lacZ$\alpha$ |
| **L2649** | agcaactgattcagccgtaaccagaaaaaagg |  |
| **L2650** | ggttacggctgaatcagttgctgcgtcgg | PCR to amplify fragment down of *cbsFHCEAP-fct* to clone in pre112-lacZ$\alpha$ |
| **L2651** | gaCTCTAGAGGATCCCCGGGTAattaccagttacccgcctcg |  |
| **L2678** | cgccagggttttcccagtcacgggcatccctgtcacaattt | PCR to amplify fragment up of *solABCDEFGHIJKL* to clone in pre112-lacZ$\alpha$ |
| **L2679** | accagtgttgccattacttagccaacgaagattacattg |  |
| **L2680** | gctaagtaatggcaacactggttggacaaaatat | PCR to amplify fragment down of *solABCDEFGHIJKL* to clone in pre112-lacZ$\alpha$ |
| **L2681** | gaCTCTAGAGGATCCCCGGGTAttaggctgtatcgggctg |  |
| **L762** | GTTATTGGTGCCCTTAAACG | Oligonucleotides used to verify insertion of fragment in pRE112-lacZ$\alpha$ and used for Sanger sequencing |
| **L763** | GCATCCAACGCCATTCATGG |  |
| **L707** | agcggataacaatttcacacagga | Oligonucleotides used to verify insertion of fragment in pSEVA421 and used for Sanger sequencing |
| **L33** | TTAATGCGCCGCTACAGGGCG |  |
| **L2480** | gcagacctttgcggaattgc | Oligonucleotides used to verify deletion of *prtBCA* |
| **L2481** | cattcaggtgctggatgtgc |  |
| **L2658** | ttccaccgggtcattccggg | Oligonucleotides used to verify deletion of *pelBCZ* |
| **L2659** | ctgaaaaccggcgaaatgcc |  |
| **L2660** | gtcaatcaacagatgcatggcg | Oligonucleotides used to verify deletion of *cbsF* |
| **L2661** | gcaaaaccaagcctggatcc |  |
| **L2662** | gcaaaacattcagtggcgtgg | Oligonucleotides used to verify deletion of *cbsFHCEAP-fct* with L2661 |
| **L2682** | ctaacagtaaaatcattggagcgct | Oligonucleotides used to verify deletion of *solABCDEFGHIJKL* |
| **L2683** | ctctgagtatatggcaagtccct |  |
| **L1833** | CTGCATGaattcccgggagagctcgccgatatcacatggcag | PCR to amplify fragment up of *budAB* to clone in pre112 |
| **L1834** | cagttggctgatcatatccattcccttctgacga |  |
| **L1835** | gaatggatatgatcagccaactgatctgagc | PCR to amplify fragment down of *budAB* to clone in pre112 |
| **L1836** | tcccaagcttcttctagaggtacccaggtgcggaacgcggcgatt |  |
| **L2062** | TGCATGaattcccgggagagctcatgcgctgatgaatcgc | PCR to amplify fragment up of *budR* to clone in pre112 |
| **L2063** | gatcaggcgcttaattccatatattttcgatctccaaatgg |  |
| **L2064** | atatggaattaagcgcctgatcgttctca | PCR to amplify fragment down of *budR* to clone in pre112 |
| **L2065** | cccaagcttcttctagaggtaccataactctccccgataacaaatcaac |  |
| **L2066** | TGCATGaattcccgggagagctcaccaccgataatcagattgtctgg | PCR to amplify fragment up of *budC* to clone in pre112 |
| **L2067** | tatcagttaaactgtttcatggtcttttcctgt |  |
| **L2068** | ccatgaaacagtttaactgatatcgtatttaactgatatcgtattcag | PCR to amplify fragment down of *budC* to clone in pre112 |
| **L2069** | cccaagcttcttctagaggtaccggacacgatgtgtagcgg |  |
| **L1874** | ccgttgggcgcgttctataa | Oligonucleotides used to verify deletion of *budAB* |
| **L1875** | cttcaaactcctcgccgatat |  |
| **L2101** | aggactacggaaaccgataacg | Oligonucleotides used to verify deletion of *budR* |
| **L2102** | gcttaagttcctctctttgcgc |  |
| **L2103** | aatgcgttcacgatcaaaacg | Oligonucleotides used to verify deletion of *budC* |
| **L2104** | ccgccatttcgtataaccgatc |  |
